# Illusions of Alignment Between Large Language Models and Brains Emerge From Fragile Methods and Overlooked Confounds

**DOI:** 10.1101/2025.03.09.642245

**Authors:** Nima Hadidi, Ebrahim Feghhi, Bryan H. Song, Idan A. Blank, Jonathan C. Kao

**Author notes:** Equal contribution, list order is random (Nima Hadidi, Ebrahim Feghhi). Co-senior authors (Idan A. Blank, Jonathan C. Kao).

## Abstract

Emerging research seeks to draw neuroscientific insights from the neural predictivity of large language models (LLMs). However, as results continue to be generated at a rapid pace, there is a growing need for large-scale assessments of their robustness. Here, we analyze a wide range of models, methodological approaches, and neural datasets. We find that some methodological approaches, particularly the use of shuffled train-test splits, have led to many impactful yet unreliable findings, and that the method by which activations are extracted from LLMs can bias results to favor particular model classes. Moreover, we find that confounding variables, particularly positional signals and word rate, perform competitively with trained LLMs and fully account for the neural predictivity of untrained LLMs. In summary, our results suggest that theoretically interesting connections between LLMs and brains on three neural datasets are driven largely by fragile methodologies and overlooked confounds.

## 1 Introduction

In recent years, large language models (LLMs) have profoundly transformed the world, demonstrating an astonishing ability to interact with humans through natural language. This raises a fundamental question: do LLMs process language in a human-like manner? One way researchers have sought to investigate this question is through the neural encoding framework, in which an LLM’s internal representations of linguistic stimuli are used to predict brain responses to the same stimuli. Across multiple studies, results have been strikingly consistent—LLM-derived representations are highly effective at predicting brain responses [Toneva and Wehbe, 2019, Schrimpf et al., 2021, Goldstein et al., 2022, Caucheteux and King, 2022, Antonello et al., 2023]. An emerging research program now seeks to understand what properties of LLMs drive this neural predictivity.

In the most cited of these studies, Schrimpf et al. [2021] evaluated the neural predictivity of 43 models—including transformer-based LLMs, smaller recurrent neural network (RNN) models, and static (i.e. non-contextual) word embeddings—across three neural datasets and reported three main results. First, LLMs trained to predict the next word, specifically *GPT2XL* Radford et al. [2019], provided the best neural predictivity across all 43 models, accounting for 100% of the explainable neural variance in one neural dataset (i.e. taking into account noise inherent in the data). Second, the neural predictivity of models was positively correlated with one model property in particular, their next-word prediction ability. These two findings were interpreted as “computationally explicit evidence that predictive processing fundamentally shapes the language comprehension mechanisms in the human brain” [Schrimpf et al., 2021]. Third, untrained (i.e. randomly initialized) transformer-based LLMs demonstrated surprisingly high neural predictivity relative to their trained counterparts, which was interpreted as evidence that the transformer architecture may play a role in biasing computations to be more brain-like. Taken together, these findings paint a picture in which the artificial intelligence community is “rapidly converging on architectures that might capture key aspects of language processing in the human mind and brain” [Schrimpf et al., 2021].

Schrimpf et al. [2021] laid the groundwork for several follow-up studies which viewed LLMs not only as predictive tools, but also as candidate explanatory models of biological language processing. For example, Hosseini et al. [2024a] demonstrated that LLMs achieve high neural predictivity even when the scale of natural language data they are trained on mirrors the developmental language exposure of humans. Aw et al. [2024] highlighted the effects of instruction tuning on LLMs, revealing that this process enhances both their alignment with neural data and their integration of world knowledge. AlKhamissi et al. [2024] further investigated the neural predictivity of untrained transformer-based LLMs, attributing it to tokenization strategy and multi-headed attention. The authors then constructed a model based on these two components that displayed high neural and behavioral predictivity, leading them to “conceptualize language processing in the human brain as an untrained feature encoder providing representations to a downstream trainable decoder that produces language output”.

While these studies represent the growing enthusiasm for using LLMs to gain neuroscientific insights, caution is warranted. LLMs are a relatively recent technology, and the field is still in the process of establishing robust methodologies to relate their representations to brain responses. Moreover, the excitement surrounding LLMs creates an incentive to identify theoretically compelling parallels between artificial and biological language processing, sometimes at the risk of overlooking simpler confounds that may drive these mappings [Bowers et al., 2022]. In this study, we empirically demonstrate these issues by showing that results from Schrimpf et al. [2021] depend on fragile methods and can be accounted for by simple confounds. Importantly, the issues we highlight have direct implications for a large set of follow-up LLM-to-brain mapping studies as well [Oota et al., 2022, Aw et al., 2024, Hosseini et al., 2024a, Kauf et al., 2024, Hosseini et al., 2024b, Mischler et al., 2024, AlKhamissi et al., 2024]. Through this work, we hope to provide a necessary course correction against drawing excessive correspondences between LLMs and brains, and encourage future research which critically evaluates parallels between artificial intelligence and neuroscience.

## 2 Results

We evaluated the neural predictivity of several models on three neural datasets: (1) *Pereira2018* (fMRI - passages) [Pereira et al., 2018], (2) *Fedorenko2016* (ECoG - sentences) [Fedorenko et al., 2016], and (3) *Blank2014* (fMRI - stories) [Blank et al., 2014]. In *Pereira2018*, n = 10 participants read short passages presented one sentence at a time, and a single fMRI volume (TR) was acquired after presentation of each sentence. *Pereira2018* consists of two sub-experiments, and we combined results across both experiments following Schrimpf et al. [2021]. In *Fedorenko2016*, ECoG recordings were made while *n* = 5 participants read 52 sentences, presented word-by-word. In *Blank2014*, fMRI signals were acquired while *n* = 5 participants listened to 8 stories. We focus all analyses on language-selective voxels (*Pereira2018*) / electrodes (*Fedorenko2016*) / fROIs (*Blank2014*). For additional details on these three neural datasets, see 4.1.

When evaluating the neural predictivity of a model, we first passed as input to the model the same natural language stimuli that the participants received. We then extracted model activations (4.3), and trained linear regressions to predict brain responses from each layer of the model. To assess a model’s neural predictivity, we used nested cross-validation (4.9). In each outer fold, the subset of brain responses and model activations used for fitting the regression is termed the train set, and the held-out subset of brain responses and model activations used to evaluate the regression is termed the test set. For models with multiple layers, we focus our analyses on the layer which achieves the highest neural predictivity on the test set (averaged across outer folds). (4.10).

### 2.1 LLM neural predictivity values generated with shuffled splits are not reliable

#### 2.1.1 A trivial model of temporal autocorrelation achieves higher neural predictivity than *GPT2XL* when using shuffled splits

The majority of studies evaluating the neural predictivity of language models have employed contiguous train-test splits [e.g., Huth et al., 2016, Antonello et al., 2023, Antonello and Huth, 2024, Caucheteux et al., 2021, Caucheteux and King, 2022, Caucheteux et al., 2023], where a temporally contiguous chunk of brain responses (and the associated model activations) are held-out for testing. By contrast, several studies using the neural datasets we study here [AlKhamissi et al., 2024, Aw et al., 2024, Kauf et al., 2024, Oota et al., 2022, Hosseini et al., 2024a,b], most notably Schrimpf et al. [2021], used shuffled splits, where samples are arbitrarily placed into the test set without regard for temporal adjacency. To provide an example with the *Pereira2018* dataset, contiguous splits imply that entire passages are held out for testing, whereas shuffled splits allow that some sentences from a given passage are placed in the train set, and other sentences from *the same passage* are placed in the test set.

Shuffled splits are known to be problematic in the field of language neural encoding because brain responses are temporally autocorrelated for reasons that are not entirely stimulus-driven [Zada et al., 2024]. Due to temporal autocorrelation in brain responses, a model can achieve high neural predictivity purely because it represents nearby time points similarly when using shuffled splits. To demonstrate this, we constructed a trivial model which operates only on the principle that stimuli nearby in time should be assigned similar representations. We term this model the Orthogonal Autocorrelated Sequences Model (*OASM*) because the activations of *OASM* are orthogonal for distinct passages/sentences/stories, and autocorrelated for stimuli within a passage/sentence/story (4.4).

We compared the neural predictivity of *OASM* to that of *GPT2XL* when using largely the same methodological choices implemented in Schrimpf et al. [2021]. These methodological choices were fitting the mapping between model activations and brain responses with ordinary least squares (OLS) regression (4.7) and extracting activations from *GPT2XL* using the last token of the stimuli, which we term *GPT2XL-LAST* (4.3). We performed a one-sided Wilcoxon signed-rank test across participants to determine if *GPT2XL* neural predictivity was significantly higher than that of *OASM* with *α* = 0.05 (4.12). Under these settings, *OASM* predicted brain responses qualitatively on-par with or better than *GPT2XL-LAST* across all three datasets, and there was no case where *GPT2XL-LAST* predicted brain responses significantly better than *OASM* (Figure 1a, top left panel). To evaluate the robustness of this finding, we compared the performance of *OASM* to *GPT2XL* when using two other activation extraction methods: taking the mean across tokens (mean pooling, referred to as *GPT2XL-MEAN*), and taking the sum across tokens (sum pooling, referred to as *GPT2XL-SUM*). We included mean pooling because it was used in Kauf et al. [2024], and sum pooling because it is somewhat analogous to the process of convolving the hemodynamic response function (HRF) with model activations (4.3). Results were consistent, with *OASM* qualitatively achieving the same or higher neural predictivity than *GPT2XL-MEAN* and *GPT2XL-SUM* across all three neural datasets.

**Figure 1:**
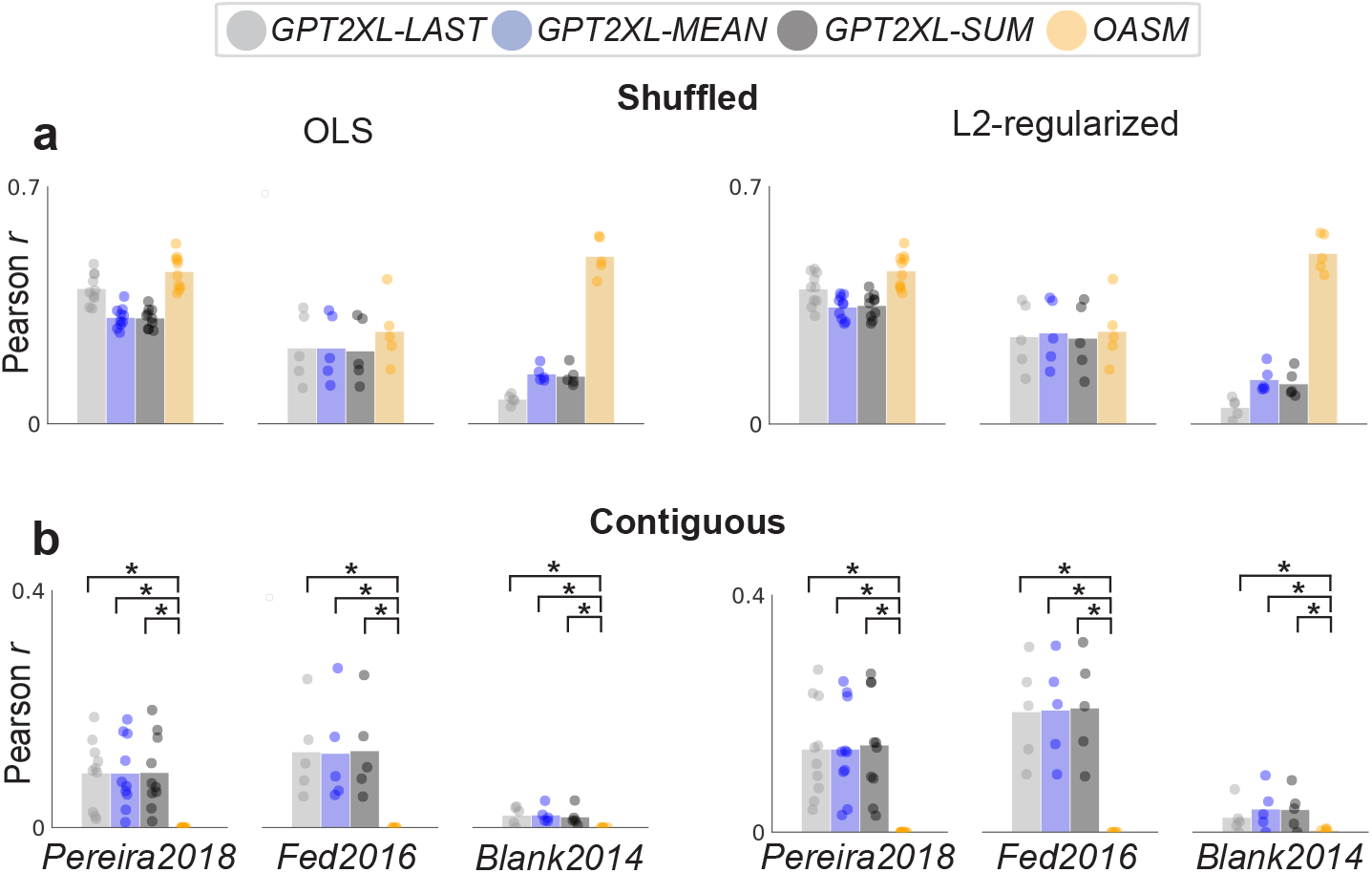
**a)** Neural predictivity of *OASM* and *GPT2XL* across three different activation extraction methods: last token pooling (*-LAST)*, mean pooling (*-MEAN)*, and sum pooling (*-SUM*). Each dot shows the mean predictivity across voxels/electrodes/fROIs for a given participant, and the bar displays the mean value across participants. Left side displays neural predictivity when using OLS regression, right side shows L2-regularized regression. **b)** Same as **(a)** but when using contiguous splits. * denotes that *GPT2XL* performs significantly better (*α* = 0.05) than *OASM* across participants using a one-sided Wilcoxon signed-rank test.

We hypothesized that the gap in neural predictivity between *GPT2XL* and *OASM* was partially due to the use of OLS regression, which does not use regularization. This is because regularization tends to help higher dimensional models, and a given layer of *GPT2XL* is higher dimensional than *OASM*. We found that while using L2-regularized regression reduced the gap between *OASM* and *GPT2XL* on *Pereira2018, GPT2XL* still did not explain significantly more neural variance than *OASM* on any of the three neural datasets (Figure 1a, top right panel).

When switching to contiguous splits, the neural predictivity of *OASM* was 0, which is expected given that *OASM* represents distinct passages/sentences/stories orthogonally (Figure 1b). The neural predictivity of *GPT2XL* decreased moderately, but remained significantly above that of *OASM*. We also found that L2-regularized regression led to higher neural predictivity for all *GPT2XL* activation extraction variants relative to OLS regression when using contiguous splits as well. For this reason, and because L2-regularized regression is the standard in the field [Huth et al., 2016, Antonello et al., 2023, Antonello and Huth, 2024, Caucheteux et al., 2021, Caucheteux and King, 2022, Caucheteux et al., 2023, Goldstein et al., 2024, 2022], we used L2-regularized regression for the remainder of our analyses.

#### 2.1.2 *OASM* accounts for the majority of the neural predictivity of *GPT2XL*

While *OASM* explains similar or more neural variance than *GPT2XL* when using shuffled splits, it remains unclear how much *OASM* accounts for the same neural variance that *GPT2XL* explains. In other words, to what extent is the neural predictivity of *GPT2XL* on shuffled splits attributable to the fact that it simply represents nearby timepoints similarly?

We addressed this question in three ways. First, we examined whether the neural predictivities of *OASM* and *GPT2XL* were positively correlated with one another across voxels/electrodes/fROIs. Second, we more directly addressed this question by performing a variance partitioning analysis at the participant-level using *R*^2^. In brief, this variance partitioning analysis first quantified the neural predictivity of a model which combines activations from both *OASM* and *GPT2XL* (*OASM+GPT2XL*). We then quantified the fraction of the neural variance that *GPT2XL* explains that is not explained by *OASM* by subtracting the neural predictivity of *OASM* from that of *OASM+GPT2XL*, and dividing this difference by the neural predictivity of *GPT2XL* alone. Finally, we subtract this fraction from 1 and convert it to a percentage to quantify the neural variance explained by *GPT2XL* that is also explained by *OASM*. This metric is termed Ω_*GPT2XL*_(*OASM*), or simply Ω (4.11). Third, because Ω is defined at the participant level, we use a one-sided paired t-test across square error values to determine the percentage of voxels/electrodes/fROIs where *OASM+GPT2XL* explained significantly more neural variance than *OASM* alone. We applied FDR correction for each participant separately for this analysis [Benjamini and Hochberg, 1995] (4.12).

Across all three datasets, neural predictivity across voxels/electrodes/fROIs was correlated between *OASM* and *GPT2XL* (Figure 2a, b). Furthermore, *OASM* accounted for over 80% of the neural variance that *GPT2XL* explains in *Pereira2018*, over 50% in *Fedorenko2016*, and nearly 100% in *Blank2014* (Figure 2c). *OASM+GPT2XL* explained significant neural variance over *OASM* in around 20% of voxels, 30% of electrodes, and 0 fROIs (Figure 2d). Overall, these results show that, when using shuffled splits, the majority of the neural predictivity of *GPT2XL* is confounded with a model that simply represents nearby time points similarly.

**Figure 2:**
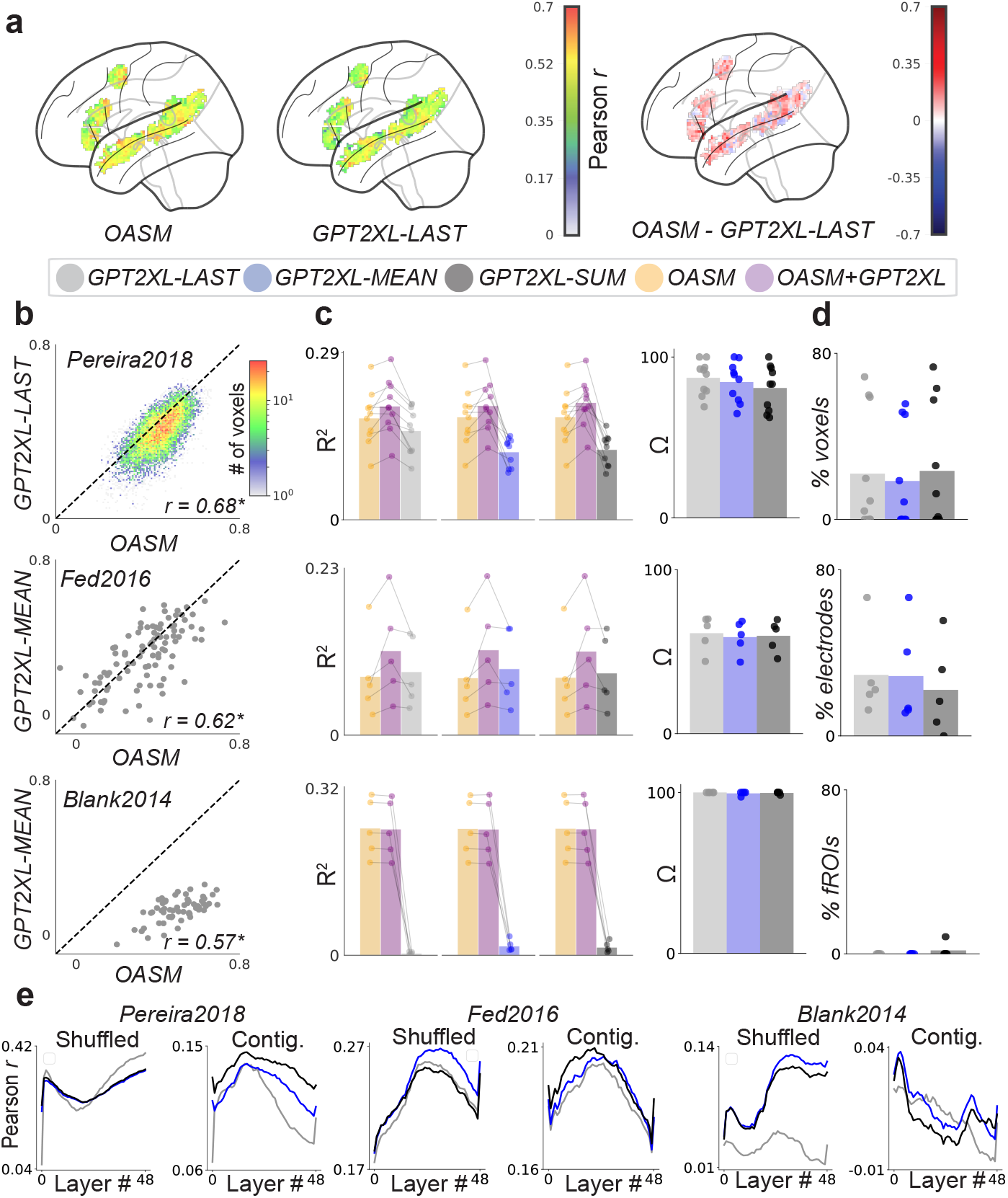
**a)** Glass brain plots showing Pearson *r* values across the left hemispheric language network in *Pereira2018*. Plots are generated by averaging Pearson *r* values within the language network across all participants. **b-d)** Top row is *Pereira2018*, middle row is *Fedorenko2016*, and bottom row is *Blank2014*. **b)** 2*D* histogram or scatter-plot comparing Pearson *r* values for *OASM* and *GPT2XL* at the voxel/electrode/fROI level. Bottom right corner shows the correlation in neural predictivity between the two models, with * indicating *p <* 0.05. **c)** Left-side shows the neural predictivity of *OASM, OASM+GPT2XL*, and *GPT2XL* in *R*^2^. Lines connect dots from the same participant. These neural predictivity values are used to compute the percentage of *GPT2XL* neural predictivity that is accounted for by *OASM*, or Ω_GPT2XL_(*OASM*), displayed on the right side. **d)** Percentage of voxels/electrodes/fROIs where *OASM+GPT2XL* explains significantly more neural variance than *OASM* alone after FDR correction within participant. **e)** Across layer neural predictivity of *GPT2XL* when using shuffled and contiguous splits. For all bar plots, each dot represents a participant, and each bar shows the mean across participants. Results in panels **(a)** and **(b)** are only shown for the best activation extraction variant of *GPT2XL*.

#### 2.1.3 Across-layer neural predictivity patterns between shuffled and contiguous splits are anti-correlated

In the previous analyses, we showed that shuffled splits are not a reliable method to assess neural predictivity. It is therefore important to assess to what extent switching from shuffled splits to contiguous splits impacts LLM-to-brain mapping findings. To begin answering this question, we examined the pattern of neural predictivity across the layers of *GPT2XL* when using shuffled vs. contiguous splits. The across-layer pattern of neural predictivity is important because studies which employed shuffled splits using these datasets exclusively focus their analyses on either the LLM layer which achieves the highest neural predictivity [Schrimpf et al., 2021, Hosseini et al., 2024a, Kauf et al., 2024, Aw et al., 2024] or the last LLM layer [Oota et al., 2022].

We found that in *Pereira2018*, the neural predictivity across layers (*n* = 48) was highly anti-correlated between shuffled and contiguous splits: *r* = *−* 0.60, *−* 0.86, and *−* 0.85 (all *p <* 0.05) for *LAST, MEAN*, and *SUM* (Figure 2e). When using shuffled splits, the early and late layers of *GPT2XL* achieved the highest neural predictivity, whereas when using contiguous splits the intermediate layers achieved the best neural predictivity. Because we selected the best layer of *GPT2XL* for each of the two experiments in *Pereira2018* separately, we also show that these same trends hold when evaluating across-layer patterns within each experiment separately (Extended Data Figure 1b). In *Fedorenko2016*, the across-layer trends were more similar between shuffled and contiguous splits (*r* = 0.61, 0.57, 0.52; all *p <* 0.05), with later intermediate layers generally performing the best on shuffled splits and early intermediate layers performing the best on contiguous splits (Figure 2e). Finally, in *Blank2014*, the across-layer performance was also anti-correlated between shuffled and contiguous splits for *GPT2XL-MEAN* and *GPT2XL-SUM* (*r* = 0.46, *−* 0.65, *−* 0.44; all *p <* 0.05); later intermediate layers performed the best when using shuffled splits, whereas the early layers performed the best when using contiguous splits (Figure 2e). We thus show that on 2 out of the 3 neural datasets, the pattern of neural predictivity across layers flips between shuffled and contiguous splits, providing a clear example of how shuffled splits can impact findings on LLM-to-brain mapping studies.

### 2.2 Reevaluating Model Comparisons and Methodological Biases

#### 2.2.1 Key Model Comparison Results Do Not Replicate with Contiguous Splits

Building on our observation that shuffled splits can alter across-layer neural predictivity trends, we next asked whether key model comparison analyses from Schrimpf et al. [2021] hold under contiguous splits. Of the models evaluated by Schrimpf et al. [2021], we use 28 of the 29 bidirectional transformer models, 6 of the 9 unidirectional transformer models (excluding those that were previously mislabeled as bidirectional, see 4.2), and 2 of the 3 static word embedding models. In addition to the models evaluated in Schrimpf et al. [2021], we evaluate 4 recent unidirectional transformer models (*Llama-3.2* family) and 4 recent unidirectional RNNs (*RWKV-4* family). Previously, Schrimpf et al. [2021] reported that unidirectional transformer models (specifically those from the *GPT2* family) predicted neural responses uniquely well compared to both bidirectional transformers and recurrent models, leading them to infer that unidirectional transformers were uniquely “brain-like”. However, these findings relied on shuffled splits and last-token activation extraction, which may have introduced biases.

Using contiguous splits and the best-performing activation extraction method for each model, we do not find that unidirectional transformers consistently achieve higher neural predictivity scores than bidirectional transformers on any dataset, nor do we find that they consistently outperform unidirectional RNNs (Figure 3a). We do replicate the finding from Schrimpf et al. [2021] that static word embedding models consistently exhibit lower neural predictivity than all contextual models (i.e. LLMs) across all three datasets.

**Figure 3:**
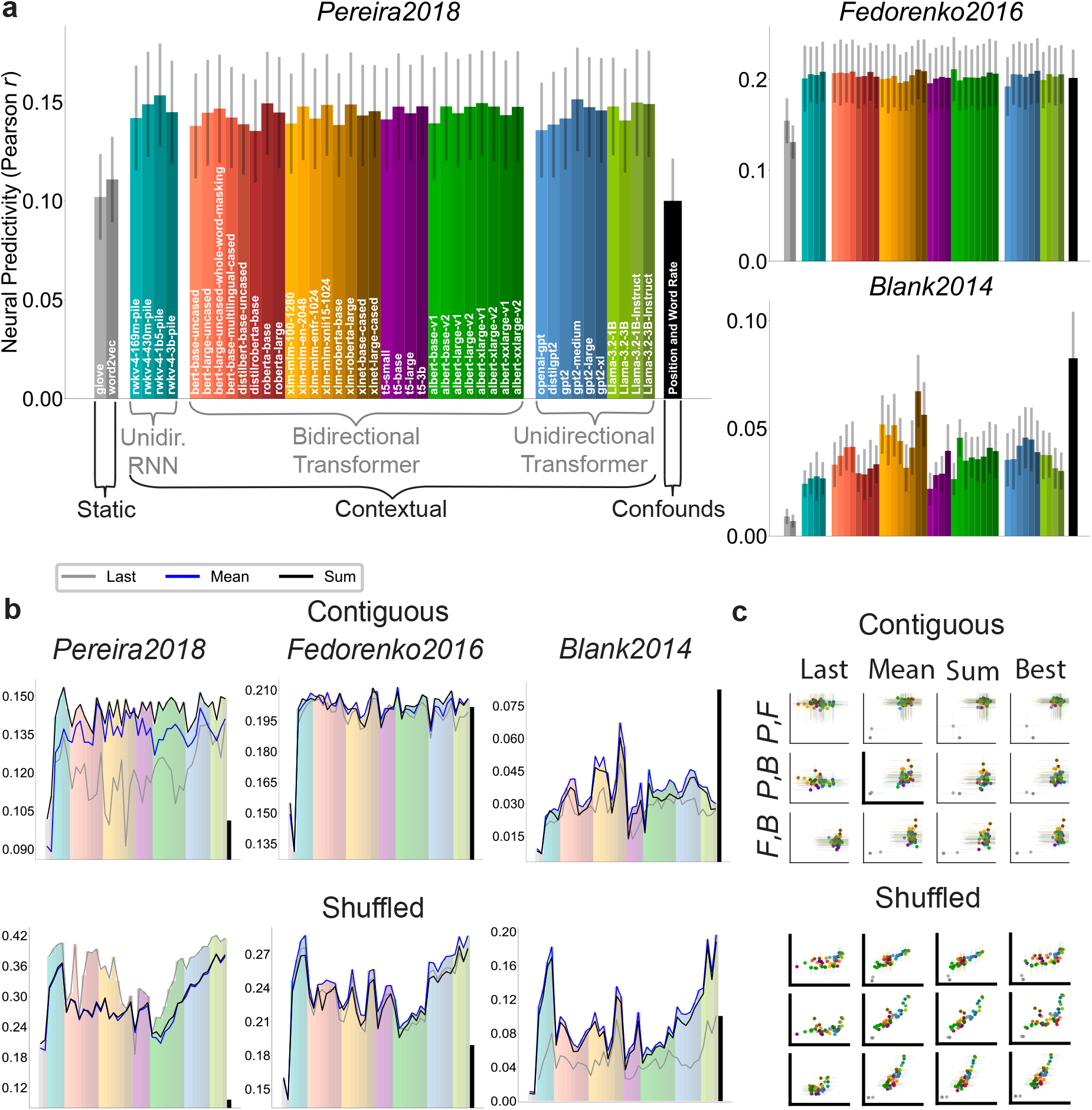
(a) Neural predictivity shown across a wide array of models when using contiguous splits and the best-performing activation extraction method for each model. **(b)** Lines represent neural predictivity for each activation extraction method. Colored bars depict best-performing activation extraction method. Top row shows results when using contiguous splits; bottom row shows shuffled splits. Note that the y-axes for these plots do not necessarily begin at 0. **(c)** Correlation in neural predictivity between datasets. Scatter plots depicting the relation in neural predictivity trends across models between each pair of datasets. *P, F*, and *B* refer to *Pereira2018, Fedorenko2016*, and *Blank2014*, respectively. Rows are labeled with the datasets plotted along the *x* and *y* axes, respectively. Columns correspond to different activation extraction methods. Axes are bolded if the Pearson correlation is significant (*p<0.05*) when static embedding models are excluded. All correlations are significant when static embedding models are included. Note that neural predictivity scores are not reported for static embedding models with the last-token method, as this method is ill-suited for non-contextual models.

However, the gap in predictivity between contextual models and static embeddings is not necessarily caused by contextual processing of linguistic content; other factors might explain this gap. In particular, we introduce a simple baseline, the Position and Word Rate (*PWR*) model, which encodes only positional signals and word rate information (4.5). While *PWR* matches static models in the *Pereira2018* dataset, it rivals contextual models in the *Fedorenko2016* dataset and outperforms them in the *Blank2014* dataset, revealing the potential impact of simple confounds in these datasets.

We next examined how the choice of activation extraction method can influence model comparisons. When examining the neural predictivity of each model under each activation extraction method (Fig. 3b top, per-participant results shown in Extended Data Fig. 4), we find that the last-token method biases the results in the *Pereira2018* dataset against bidirectional models. To assess significance, we performed a one-sided Wilcoxon signed-rank tests between each unidirectional transformer model and each bidirectional transformer model (Extended Data Fig. 5). Whereas 51.1% of comparisons showed significantly greater neural predictivity for unidirectional transformer models when using the commonly used last-token activation extraction method, less than 10% of comparisons were significant when using either mean pooling, sum pooling, or the best activation extraction method per model. Hence, our results highlight that even under contiguous splits, the choice of activation extraction method can greatly affect model comparison results. This is typically underappreciated in the field, as the majority of LLM-to-brain mapping findings depend on a single activation extraction method.

**Figure 4:**
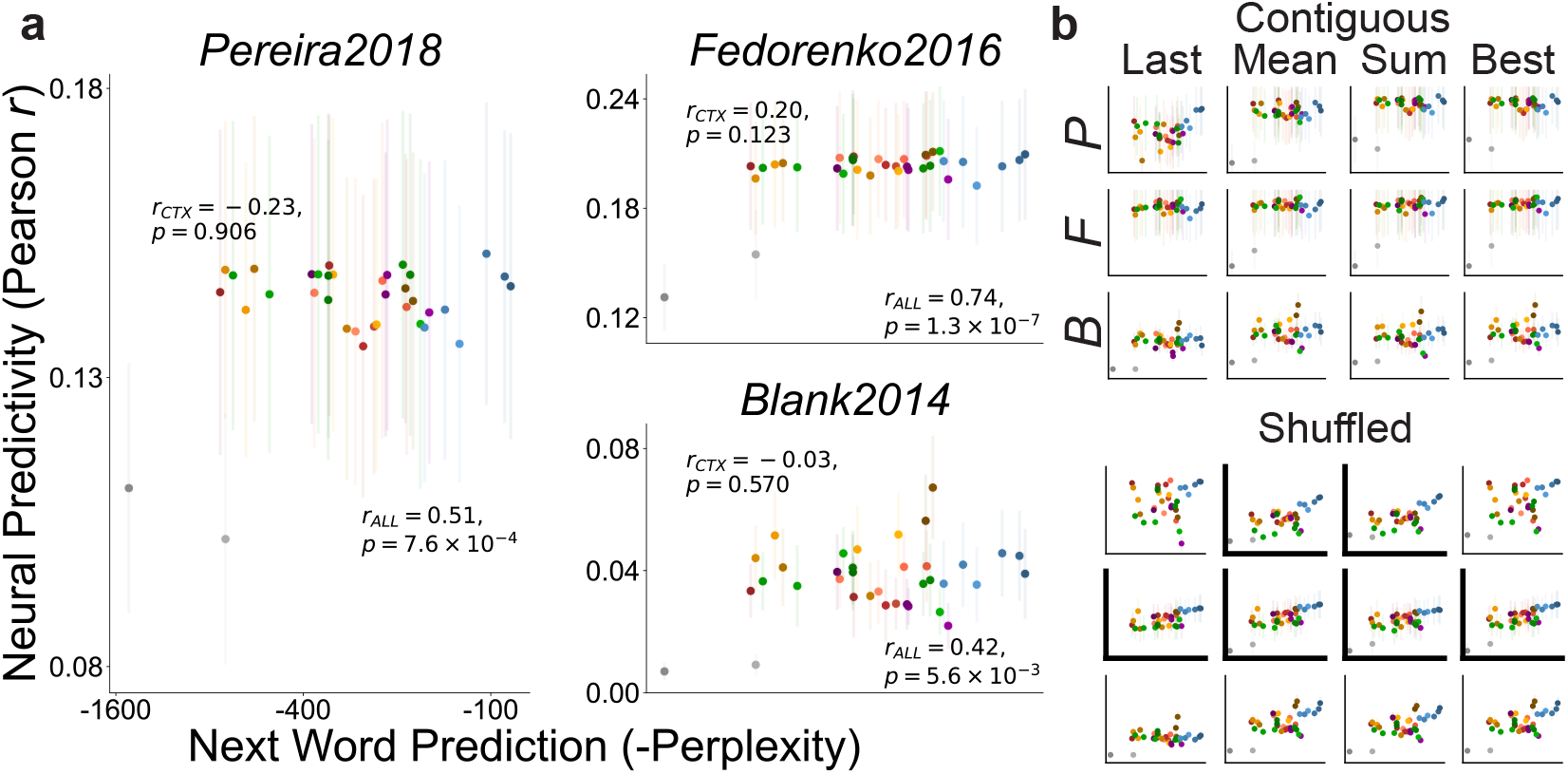
**(a)** Scatter plots depict the relation between next-word prediction and neural predictivity on each dataset when using contiguous splits and the best-performing activation extraction method for each model. Pearson correlations and p-values are given when including all models (*r*_*ALL*_) and when only including contextual models (*r*_*CTX*_). **(b)** Scatter plots depict the relation between next-word prediction and neural predictivity across models for each dataset (rows) with each activation extraction method (columns) with contiguous (top) and shuffled (bottom) splits. Axes are bolded when r_*CTX*_ is significant (p<0.05). r_*ALL*_ is significant in all cases where applicable; r_*ALL*_ is not defined when using the last-token method as static embedding model scores are omitted in this case.

**Figure 5:**
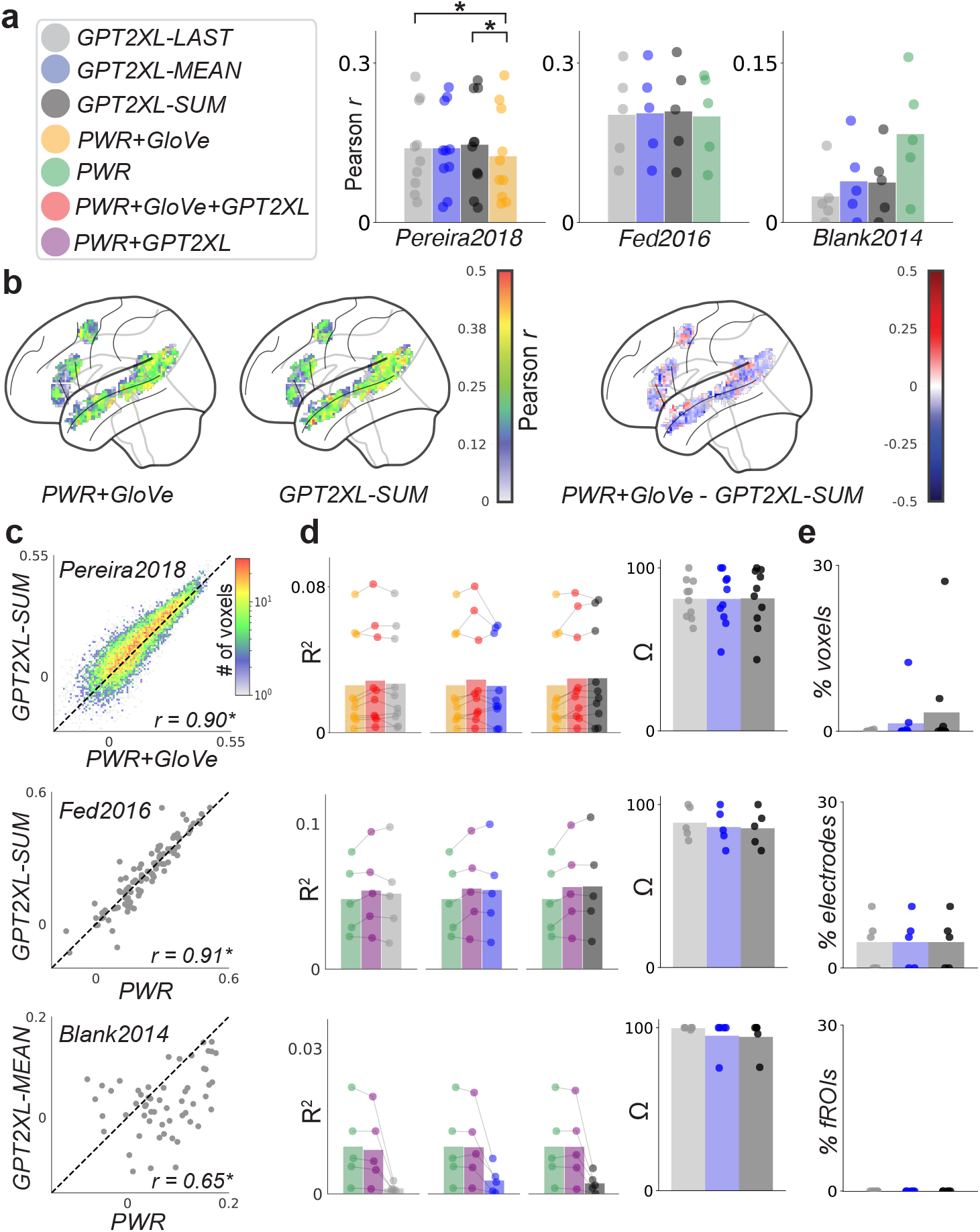
**a)** Neural predictivity of different models, in Pearson *r*, for each neural dataset. Asterisks indicate that *GPT2XL* neural predictivity is significantly higher than that of *PWR+GloVe* / *PWR* (*α* = 0.05) **b)** Glass brain plots showing Pearson *r* values across the left hemisphere of the language network in *Pereira2018*. Plots are generated by averaging Pearson *r* values within the language network across all participants. **c)** 2*D* histogram or scatter-plot comparing Pearson *r* values for *PWR+GloVe* / *PWR* and *GPT2XL* at the voxel/electrode/fROI level. Bottom right corner shows the correlation in neural predictivity between the two models, with * indicating *p <* 0.05. **d)** Left side shows the neural predictivity of *PWR+GloVe* / *PWR, PWR+GloVe+GPT2XL* / *PWR+GPT2XL*, and *GPT2XL* with *R*^2^. Lines connect dots from the same participant. These neural predictivity values are used to compute percentage of *GPT2XL* neural predictivity that is accounted for by *PWR+GloVe / PWR* displayed on the right side. **e)** Percentage of voxels/electrodes/fROIs where *PWR+GloVe+GPT2XL* / *PWR+GPT2XL* explains more neural variance than *PWR+GloVe* / *PWR* alone. For all bar plots, each dot shows values for a given participant, and bars show means across participants. Results in panels **(b)** and **(c)** are only shown for the best activation extraction variant of *GPT2XL*.

When reintroducing shuffled splits (Figure 3b bottom), unidirectional transformers often outperformed bidirectional transformers across datasets and activation extraction methods. Indeed, for all activation extraction methods on all datasets, a greater fraction of comparisons between unidirectional transformers and all other classes of models (bidirectional transformers, unidirectional RNNs, and static word embeddings) were significant when using shuffled splits than when using contiguous splits (Extended Data Fig. 5). This suggests that the use of shuffled splits likely drove the apparent superiority of unidirectional transformer models in Schrimpf et al. [2021].

### 2.2 Relative Performance Trends Across LLMs Are Inconsistent Between Datasets

Schrimpf et al. [2021] also reported that relative performance trends across models were correlated between datasets; we next asked whether such inter-dataset robustness persists under contiguous splits. Following their analyses, we examine the consistency of neural predictivity trends by calculating Pearson correlations across model neural predictivity scores (Pearson *r*) between datasets (Figure 3c). Using contiguous splits and the best activation extraction method for each model, significant correlations emerged when all models were included, but vanished when static word embedding models were excluded. This pattern persisted when using the same activation extraction method across all models: significant correlations were always found when including all models, but when excluding the two static word embedding models, a significant correlation was found in only 1 out of 9 cases: between *Pereira2018* and *Blank2014* when mean-pooling. Under shuffled splits, significant correlations appeared universally across datasets and activation extraction methods, regardless of whether static models were included. These findings indicate that shuffled splits in Schrimpf et al. [2021] likely exaggerated the consistency of differences in neural predictivity among LLMs across datasets.

#### 2.2.3 Correlation Between Neural Predictivity and Next-Word Prediction Is Fragile

Finally, we reassessed the most impactful result from Schrimpf et al. [2021]: the correlation between next-word prediction (quantified as negative perplexity) and neural predictivity. For these analyses, we only used the 36 models that were also used by Schrimpf et al. [2021] so that we could use the same next-word prediction metrics they had previously reported. Using contiguous splits and the best activation extraction method per model, we found significant correlations in all datasets only when static word embedding models were included. When restricted to contextual models, these correlations disappeared (Figure 4). These results were consistent across all activation extraction methods: significant correlations were only found when including static word embedding models, suggesting that the correlation merely reflects a difference between static word embedding models and contextual models. Under shuffled splits, correlations were more widespread across datasets and activation extraction methods, appearing in 6 out of 12 cases. Overall, our findings demonstrate that the correlation between neural predictivity and next-word prediction on these datasets is far less robust than previously reported.

### 2.3 Positional information, word rate, and static embeddings account for the majority of the neural predictivity of trained LLMs

Our analyses from the model comparison section 2.2 reveal that Position and Word Rate (*PWR*) as well as static word embedding models, perform surprisingly well relative to LLMs. Motivated by this finding, we sought to quantify what percentage of the neural predictivity of LLMs can be accounted for by *PWR* and static models. To address this question, we use *GloVe* and *GPT2XL* as representative examples of static and contextual models, respectively.

We first fit a joint regression with activations from both *PWR* and *GloVe* (*PWR+GloVe*) for each of the three neural datasets. *PWR+GloVe* achieved better neural predictivity than either model alone on *Pereira2018*; on *Fedorenko2016* and *Blank2014, PWR+GloVe* did not achieve better neural predictivity than *PWR* alone, and so we used *PWR* for further analyses with these datasets. Across voxels/electrodes/fROIs, the neural predictivity of *PWR+GloVe* / *PWR* was strongly correlated with the neural predictivity of *GPT2XL* (Figure 5b, c). Furthermore, *PWR+GloVe* accounted for over 85% of the neural variance explained by *GPT2XL* in *Pereira2018*, and *PWR* accounted for over 80% and nearly 100% of the neural variance *GPT2XL* explained in *Fedorenko2016* and *Blank2014*, respectively. Finally, *PWR+GLoVe+GPT2XL* explained significant neural variance over *PWR+GloVe* in only around 10% of voxels, and *PWR+GPT2XL* explained significant neural variance over *PWR* in only around 5% of electrodes and 3% of fROIs.

We replicated these key findings when using three other LLMS, *RoBERTa-Large, RWKV*, and *Llama* (Extended Data Figure 2). The one exception was that Ω values on *Blank2014* were lower with these three LLMs relative to *GPT2XL* (i.e., *PWR* accounted for a lower percentage of the neural variance explained by *GPT2XL*). However, these values are likely lower because accounting for the neural variance that an LLM explains is unstable when LLM neural predictivity is itself very low, as is the case on *Blank2014* (see 4.11 for more detail). Importantly, there are no fROIs for which any of these three LLMs explain significantly more neural variance than *PWR* on *Blank2014*.

A previously explored scientific question on *Pereira2018* is whether the mapping between LLMs and brains on *Pereira2018* reflects lexical-semantics or syntactic information [Kauf et al., 2024]. Here, we found that combining *GloVe* with *PWR* explained more neural variance than either model alone, suggesting that lexical-semantic content (which we model with *GloVe*) plays an important role in explaining LLM-to-brain mappings beyond simple confounds. To further examine this, we examined how much of the neural variance explained by *GloVe* could be accounted for by simple confounds. We found that Ω_GloVe_(*PWR*) was around 55% and that *PWR+GloVe* explained significant neural variance over *PWR* in 7.6% of voxels (Extended Data Figure 3), indicating that *GloVe* indeed explains neural variance over simple confounds. By contrast, an LLM-based model of contextual syntactic representations (*SYNTAX*) (4.6) explained much less neural variance over these simple confounds: Ω_SYNTAX_(*PWR*) was over 80%, and the *PWR+SYNTAX* model only explains significant neural variance over *PWR* in 0.00066% of voxels.

In summary, we find that a combination of positional signals, word rate, and static word embeddings predicts almost as much —or more —neural variance than *GPT2XL*. Furthermore, in all three datasets, these models account for upwards of 80% of the neural variance that *GPT2XL* explains, and these results replicate with three other LLMs. These findings suggest that even when using contiguous splits, there are relatively simple explanations underlying the neural predictivity of LLMs on these datasets.

### 2.4 Positional information and word rate fully account for the neural predictivity of Untrained GPT2XL

Having accounted for the majority of the neural variance explained by trained LLMs, we next examined whether *PWR* could account for the neural variance explained by untrained *GPT2XL* (*GPT2XLU*). Previously, the surprisingly high neural predictivity of untrained transformers led to the conclusion that the transformer architecture itself biases computations to be more brain-like Schrimpf et al. [2021], and inspired the development of a novel architecture that reportedly achieved state-of-the-art neural and behavioral alignment across several datasets AlKhamissi et al. [2024]. We hypothesized that, when evaluated on contiguous (rather than shuffled) splits, the neural predictivity of *GPT2XLU* could be fully accounted for by *PWR*.

Confirming our hypothesis, we found that *PWR* performed on par with *GPT2XLU* across all three neural datasets (Figure 6a). Furthermore, the neural predictivity of *PWR* and *GPT2XLU* was strongly correlated across voxels/electrodes/fROIs (Figure 6b,c). *PWR* accounted for essentially all (*>* 98%) of the neural variance of *GPT2XLU* across all three datasets, and there were no voxels/electrodes/fROIs where *PWR+GPT2XLU* explained significant neural variance over *PWR*. We therefore conclude that a combination of positional signals and word rate information not only explains equal or more neural variance than *GPT2XLU*, but that these simple features also explain nearly all of the neural variance that *GPT2XLU* explains.

**Figure 6:**
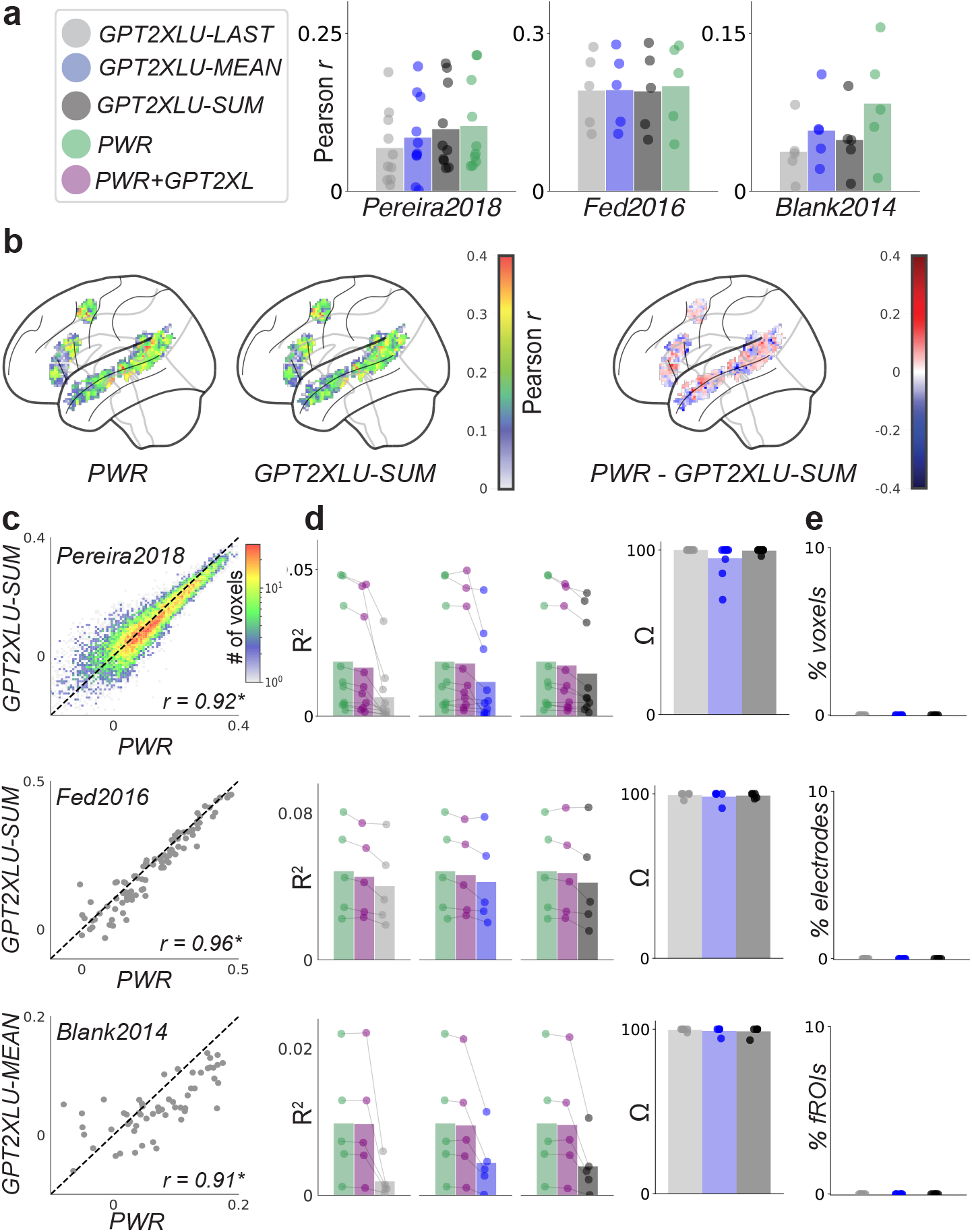
**a)** Neural predictivity of different models, in Pearson *r*, for each neural dataset. **b)** Glass brain plots showing Pearson *r* values across the left hemisphere of the language network in *Pereira2018*. Plots are generated by averaging Pearson *r* values within the language network across all participants. **c)** 2*D* histogram or scatter-plot comparing Pearson *r* values for *PWR* and *GPT2XL* at the voxel/electrode/fROI level. Bottom right corner shows the correlation in neural predictivity between the two models, with * indicating *p <* 0.05. **d)** Left side shows the neural predictivity of *PWR, PWR+GPT2XL*, and *GPT2XL* with *R*^2^. Lines connect dots from the same participant. These neural predictivity values are used to compute percentage of *GPT2XL* neural predictivity that is accounted for by *PWR* displayed on the right side. **e)** Percentage of voxels/electrodes/fROIs where *PWR+GPT2XLU* explains significantly more neural variance than *PWR* alone. For all bar plots, each dot shows values for a given participant, and bars show mean across participants. Results in panels **(b)** and **(c)** are only shown for the best activation extraction variant of *GPT2XLU*.

## Discussion

Our rigorous re-evaluation of LLM-to-brain mappings on three neural datasets offers four main contributions. First, we provide compelling evidence that shuffled splits, which were used in the most impactful study on LLM-to-brain mappings [Schrimpf et al., 2021], as well as in multiple follow-up studies [Oota et al., 2022, AlKhamissi et al., 2024, Aw et al., 2024, Hosseini et al., 2024a,b, Kauf et al., 2024, Mischler et al., 2024], are not a reliable method to assess neural predictivity. Second, when switching to standard methodological choices, namely contiguous splits and L2-regularized regression, the main results reported in Schrimpf et al. [2021] based on trained LLMs were not robust. Namely: unidirectional transformers did not exhibit better neural predictivity than other classes of LLMs; LLMs did not exhibit consistent trends in neural predictivity scores across neural datasets; and, critically, next-word prediction ability was not robustly correlated with neural predictivity.

Third, we found that positional signals and word rate (*PWR*) achieved neural predictivity on par with *GPT2XL* on two out of three neural datasets, and achieved similar or better neural predictivity relative to untrained GPT2XL (*GPT2XLU*) on all three neural datasets. Finally, a combination of *PWR* and static word embeddings (*GloVe*) accounted for over 80% of the neural variance that *GPT2XL* explains, and *PWR* alone accounts for essentially all the neural variance that *GPT2XLU* explains.

Although the majority of studies in the field use contiguous splits, rather than shuffled splits as in Schrimpf et al. [2021], we demonstrate the potential impact of shuffled splits on prior influential findings. Given that many studies that followed-up on [Schrimpf et al., 2021] drew conclusions that have not been independently replicated by other groups, our analysis suggests that all those claims warrant extensive examination. Such an examination has not been conducted to date, potentially due to two factors. First, the use of shuffled splits is often not explicitly reported. Among the eight studies we identified as employing shuffled splits, only one—Kauf et al. [2024]—clearly stated this choice; the others generally described the number of cross-validation folds without detailing their construction. Second, the methodological implications of shuffled splits have been underappreciated. For instance, Kauf et al. [2024] acknowledged that shuffled splits can “inflate” neural predictivity compared to contiguous splits, but still exclusively utilized shuffled splits for their main analyses. Our results highlight that this inflation is not uniform, resulting in markedly different patterns of neural predictivity across layers and across models, jeopardizing conclusions that follow from such layer or model comparisons.

Kauf et al. [2024] further discussed both merits and limitations of shuffled splits, with the main merit being increased semantic coverage in the training set, and the main limitation being susceptibility to temporal autocorrelation. However, their exclusive use of shuffled splits for their main analyses may inadvertently signal to the field that the merits outweigh the limitations. Our findings instead demonstrate that the issue of temporal autocorrelation is substantial, as the majority of neural variance explained by *GPT2XL* is confounded with *OASM*. Additionally, the purported benefit of increased semantic coverage with shuffled splits appears limited in the *Pereira2018* dataset, which includes multiple passages for each semantic topic. Different passages on the same topic can be assigned to training vs. test sets and, by designing our splits to leverage this feature 4.9, we demonstrate that contiguous splits can effectively balance semantic diversity and reliable estimation of model performance.

In addition to commenting on shuffled splits, our findings underscore the importance of systematically investigating multiple activation extraction methods from LLMs when fitting encoding models to fMRI datasets. In *Pereira2018*, we observe that the most commonly used approach, last-token extraction, generally performs the worst and particularly disadvantages bidirectional transformers. Differences between extraction methods do not only affect analyses that compare different LLMs (and other computational models). For example, Jain et al. [2020] demonstrated that the standard activation extraction method for narrative comprehension datasets (analogous to sum-pooling) can lead to artifacts where LLM units with the longest timescales of context integration are mapped to brain regions known to have short integration timescales, such as primary auditory cortex. More broadly, our study along with that of Jain et al. [2020] highlights a fundamental concern when interpreting LLM-to-brain mappings: the associations between model representations and brain regions are influenced by the assumptions inherent in the chosen activation extraction method.

In addition, an important finding in our study is that simple confounds, namely, positional signals and word rate, play an important role in LLM-to-brain mappings, even when using contiguous splits on these neural datasets. Most studies do not include simple confounds in the regression that maps LLM representations to neural activity, although few studies did account for such confounds. Caucheteux et al. [2021] controlled for word rate and phonological features before examining how much of the encoding performance of LLMs can be explained by syntax vs. semantics. Reddy and Wehbe [2021] iteratively accounted for simpler features, such as punctuation, before analyzing the neural predictivity of more complex syntactic models and LLM models. LeBel et al. [2021] and de Heer et al. [2017] also iteratively accounted for the neural predictivity of simpler acoustic and phonetic features before introducing more complex, semantic models. Our findings highlight the importance of the continued use of these low-level confounding predictors, especially when scientists seek to draw inferences about higher-level representations in the brain based on differences in neural predictivity between models.

The consideration of positional confounds specifically has been relatively limited to the best of our knowledge. Antonello et al. [2023] were the first to highlight this issue, identifying positional signals in the initial 100 seconds of story stimuli. To mitigate the influence of these signals, they excluded the first 100 seconds of a held-out story used in the testing set. Similarly, we observe significant positional effects at the beginnings of stories in the *Blank2014* dataset, and the neural variance predicted by *GPT2XL* is largely confounded with these signals. This finding suggests that positional confounds may be pervasive across narrative comprehension datasets. On *Pereira2018* and *Fedorenko2016*, the positional signals we model are likely confounded with aspects of the linguistic materials themselves. For instance, we find that positional signals explain roughly double the neural variance on *EXP2* relative to *EXP3* in *Pereira2018* (Extended Data Figure 2a). This difference may be due to the fact that in *EXP2* passages follow a more consistent, Wikipedia-like structure, where the first sentence defines a term, and subsequent sentences in the passage expand upon the introduced term. In *Fedorenko2016*, many sentences have similar syntactic structure, confounding linguistic content with word position. Moreover, neural activity in many electrodes increases from one word to the next in a manner that reflects the construction of meaning across words (e.g., this increase is less pronounced for lists of unrelated words [Fedorenko et al., 2016]). Hence, although we find that positional signals can explain neural responses well in these datasets, it is unclear whether the predicted brain responses reflect position-correlated linguistic content, genuine representations of position, or both.

Returning to the larger picture, our study provides an updated view regarding the “brain-likeness” of LLMs. Arguably the biggest claim about convergence between deep learning models and brains in Schrimpf et al. [2021] was that the models that best predict brain responses are precisely those that best predict the next word. Here, we find that this central finding is not robust, and depends on choice of neural datasets, models, and activation extraction methods. Results in the literature are similarly mixed. For instance, Pasquiou et al. [2022] reported a correlation between perplexity and neural predictivity within, but not across, model classes. Caucheteux and King [2022] trained 36 transformer models from scratch and found that although trained models exhibited higher neural predictivity than untrained models, neural predictivity ceased scaling with next-word prediction at some point during model training. On the other hand, **?**, Hong et al. [2024] and Bonnasse-Gahot and Pallier [2024] did report significant correlations between neural predictivity and next-word prediction. Altogether, the appearance of this correlation seems to depend somewhat unpredictably on the choice of dataset, models, and methods.

Beyond the next-word prediction abilities of LLMs, other performance metrics have also been reported to correlate with neural predictivity by studies using shuffled splits. For instance, Aw et al. [2024] reported that instruction-tuning large language models (LLMs) enhances their neural predictivity on *Pereira2018* and *Blank2014*, and showed that this increase in neural predictivity correlated with an increase in the models’ world knowledge. Mischler et al. [2024] reported that as LLMs achieve better performance on semantic benchmark tasks, their neural predictivity increases on an intracranial EEG dataset. Crucially, given that we demonstrated that the relationship between neural predictivity and one model performance metric (next-word prediction ability) can change substantially when shifting from shuffled to contiguous splits,we strongly recommend evaluating whether these other relationships are robust under contiguous splits.

A second major claim put forth in Schrimpf et al. [2021], as well as two follow-up studies Hosseini et al. [2024a], AlKhamissi et al. [2024], is that unidirectional transformers can achieve high neural predictivity when trained on little or no linguistic input, and thus these models are endowed with architectural biases that lead to “brain-like” computations. Schrimpf et al. [2021] first showed that untrained unidirectional transformers achieve surprisingly high neural predictivity on these datasets, and AlKhamissi et al. [2024] further attributed this neural predictivity to two key components: multi-headed attention and byte-pair encoding. By contrast, we show that when using contiguous splits, positional signals and word rate fully account for the neural predictivity of *GPT2XLU*. Hosseini et al. [2024a] responded to the common criticism that LLMs are not viable models of human language processing because they are trained on massive corpora of text by showing that LLMs, specifically *GPT2*-style models, can achieve high neural predictivity on *Pereira2018* when trained on developmentally realistic amounts of data. Importantly, when using shuffled splits, we find that a model that is trained on no text and which only knows passage/sentence/story boundaries (*OASM*) achieves comparable neural predictivity as fully trained *GPT2XL*, suggesting that such results are not indicative of a model’s developmental plausibility.

Beyond accounting for the neural predictivity of untrained models, positional information and word rate, along with static word embeddings, accounted for the majority of neural variance across four classes of trained LLMs. This finding, along with the fact that LLMs of all classes exhibit similar neural predictivity, suggests that trained LLMs are all mapping to similar features on these neural datasets. Interestingly, Hosseini et al. [2024b] also reinterpreted the *Pereira2018* results originally reported by [Schrimpf et al., 2021], arguing that a common set of “universal” representations across model classes are responsible for driving neural predictivity. Although their analyses are compromised by the use of shuffled splits, we are in agreement that on these datasets different classes of LLMs predict shared neural variance.

A major theme throughout our results is that the neural predictivity of LLMs can largely be explained by simpler features, making it difficult to isolate what linguistic features the brain responses reflect. For instance, we found that the neural variance predicted by contextual syntactic representations could be almost entirely accounted for by positional signals and word rate information. These simple confounds also accounted for the majority of the neural variance that a lexical-semantic model, *GloVe*, explained. In previous studies using shuffled splits, Kauf et al. [2024] claimed through a series of linguistic perturbations that lexical-semantic information, not syntactic structure, accounted for the majority of *GPT2XL* neural predictivity on *Pereira2018*, while Oota et al. [2022] claimed that fine-tuning *BERT* on syntax-related natural language processing tasks (NLP) led to greater improvements in neural predictivity relative to other NLP tasks (also when using shuffled splits). However, our results suggest that even when using contiguous splits, the *Pereira2018* dataset is not well-suited for isolating the linguistic features that drive LLM neural predictivity as these features are highly confounded with much simpler explanations.

There are three important limitations to consider when interpreting the results of our study. First, we used only three neural datasets; whereas most prior studies have used fewer datasets, the ones used here are nonetheless limited in scope. We focus on these datasets because the aim of our study was to reanalyze results from Schrimpf et al. [2021], and because they are the most commonly used datasets for neural encoding studies with localization of the language network. Future work analyzing methodological robustness and confounds should incorporate larger and more diverse datasets —including longer naturalistic narratives (e.g. LeBel et al. [2023]) —to enhance generalizability. Second, although we applied banded ridge regression to mitigate cases where stacking models together (e.g. *PWR+GloVe+GPT2XL*) decreases neural predictivity relative to a subset of those models, our procedure might still be biased against regressions with many models due to noise in neural data and low sample sizes. Finally, our Ω metric can become unstable when the neural variance that an LLM explains is near 0 (as in *Blank2014*), because that variance is in the denominator of this metric. These limitations underscore the need for further research to validate and extend our findings.

Our study joins an emerging literature in cautioning against over-interpreting a model’s high neural predictivity as evidence that the model shares theoretically interesting properties with brains [Schaeffer et al., 2022, Bowers et al., 2022, 2023, Guest and Martin, 2023, Antonello and Huth, 2024]. Large-scale analyses like ours, which incorporate diverse datasets, models, and methodologies, are essential for critically evaluating whether any specific model attribute is reliably associated with neural predictivity. Such analyses have proven similarly insightful in the visual domain, where they have likewise highlighted methodological fragility (Soni et al. [2024]) and revealed a striking similarity in the neural predictivity of deep learning models regardless of their training objective or architecture Conwell et al. [2024]. We hope that through more rigorous evaluations, the field will come to an accurate view of the correspondence between LLMs and brains.

## 4 Online Methods

### 4.1 Experimental data

For all three datasets, we used the same versions as used by [Schrimpf et al., 2021]. All analyses were done on language-selective voxels/electrodes/fROIs (see Supplementary for a description of functional localization).

#### Pereira 2018 (fMRI)

*Pereira2018* is composed of two experiments. Experiment 2 (EXP2) consists of 96 passages each containing 4 sentences (384 sentences total), with *n* = 9 participants. Experiment 3 (EXP3) consists of 72 passages each consisting of 3 or 4 sentences (243 sentences total), with *n* = 6 participants. Passages in each experiment were evenly divided into 24 semantic categories which were not related across experiments (4 passages per category in EXP2, and 3 passages per category in EXP3). Each sentence was presented for 4 s, and a single fMRI volume (TR) was acquired 4s after visual presentation of the sentence has ended. Unless otherwise noted, we focus our results on voxels from within the “language network” as identified in the main paper [Fedorenko et al., 2024] using a separate “localizer” task. Data from EXP2 consisted of a 384 *×* 92450 matrix (sentences *×* voxels) and data from EXP3 consisted of a 243 *×* 60100 matrix. For all analyses except for voxel-wise statistical testing (see 4.12), we analyze each voxel in each experiment separately, and then combine the results across voxels and experiments as done in [Schrimpf et al., 2021, Aw et al., 2024, Kauf et al., 2024, Hosseini et al., 2024a].

#### Fedorenko2016 (ECoG)

Participants (*n* = 5) read 52 sentences, each consisting of 8 words, one word at a time (416 words total). A total of 97 language-responsive electrodes were identified across 5 participants: 47, 8, 9, 15, and 18, for participants 1 through 5, respectively. High gamma activity was extracted from the neural recordings, and responses were temporally averaged across the full presentation time-window of each word. The entire dataset consisted of a 416 *×* 97 matrix (words *×* electrodes).

#### Blank2014 (fMRI)

*Blank2014* consisted of 5 participants listening to 8 stories from the publicly available Natural Stories Corpus [Futrell et al., 2018]. An fMRI volume was acquired every 2 s, resulting in a total of 1317 TRs across the 8 stories. fMRI BOLD signals were averaged across voxels within each functional region of interest (fROI) in the language network. There were 12 fROIs per participant (6 per hemisphere) for a total of 60 fROIs across all 5 participants, resulting in a 1317 *×* 60 matrix.

### 4.2 Language models

For the majority of our analyses, we focus on *GPT2XL* [Radford et al., 2019]. *We do so because GPT2XL* was shown to be the best-performing model on *Pereira2018*, and performed on par or better than other models in *Fedorenko2016* and *Blank2014* [Schrimpf et al., 2021]. *Additionally, it has been the main language model used in other neural encoding studies using these datasets [Hosseini et al., 2024a, Kauf et al., 2024]. GPT2* is a unidirectional transformer model, meaning that it can only attend to current and past tokens (but not future tokens). It is trained on next-token prediction. The *XL* variant has *∼* 1.5*B* parameters and 48 layers. We additionally replicate our findings in Figure 5 with three other models: *RoBERTa-Large, Llama-3.2-3B-Instruct*, and *RWKV-4-3B-Pile*.

Due to differences in methodological choices with Schrimpf et al. [2021], we also tested 45 models to re-evaluate their encoding performance, and the correlation between encoding performance and next-word prediction ability across those models. Out of the 45 models we used, 36 were also used in Schrimpf et al. [2021]. We omitted 7 models from Schrimpf et al. [2021]: 3 models were wrongly labeled as bidirectional models when they were in fact unidirectional (*Transfo-XL-WT103, XLM-CLM-EnFr-1024, CTRL*), and we replaced them with more recent models from the *Llama-3.2* family, including instruction-tuned variants; 2 recurrent models (*LSTM-LM1B* and *Skip-Thoughts*) were replaced with models from the *RWKV-4* family, which are more comparable in terms of their scale and performance on language tasks to the other transformer-based LLMs; 1 bidirectional model, *T5-11b*, was not included due to memory constraints; and 1 word embedding model, *ETM*, was not included because it was trained on 4 to 5 orders of magnitude less text than *GloVe* or *word2vec*.

### 4.3 LLM activation extraction and pooling

LLMs provide a representation for each token, which is either a sub-word or complete word. However, the brain responses in *Pereira2018* and *Blank2014* are acquired at a much slower temporal resolution, and so the token representations of the LLM need to be pooled to match the temporal resolution of the brain responses. The need for pooling also applies to *Fedorenko2016* in cases where tokens correspond to sub-words, because brain responses in *Fedorenko2016* are provided for each word. For all three neural datasets, we extract features from LLMs in three distinct ways: last token pooling, mean pooling, and sum pooling. We describe these methods in more detail for each dataset below. We use last token pooling because it was used in Schrimpf et al. [2021] and Kauf et al. [2024]; mean (average) pooling because it was included in Kauf et al. [2024]; and sum pooling because it is more analogous to the word-level impulse response model commonly used in more naturalistic neural encoding studies [Huth et al., 2016, Jain et al., 2020], and the HRF convolved with the brain responses to each word is similar to summing the model activations for each word.

#### Pereira2018

To extract the representation of a single sentence, we fed an LLM all words from the beginning of that sentence’s passage until and including the sentence itself. Because each fMRI volume was acquired at the end of a sentence, we converted LLM token-level representations to sentence-level embeddings by either taking the representation at the last token (which is always a period) of the sentence (last token pooling), taking the mean across tokens within the current sentence (mean pooling), or summing across tokens within the current sentence (sum pooling).

#### Fedorenko2016

For each word, its corresponding tokens were fed into the LLM together with all preceding tokens in the sentence as context. Because the brain response is averaged per-word, we converted LLM token-level embeddings to word embeddings by either taking the last token for multi-token words (last token pooling), averaging across tokens in multi-token words (mean pooling), or summing across tokens in multi-token words (sum pooling). We left single-token words unmodified.

#### Blank2014

Following Schrimpf et al. [2021], we used a version of the text that was binned into chunks corresponding to each TR. The last word in each chunk of text was approximately 4 s before the corresponding TR, reflecting the delay of the hemodynamic response. For each story, we fed all the chunks of text corresponding to that TR and all prior TRs into the LLM, up to a maximum context size of 512 tokens. For last token pooling, we selected the representation of the last token in the current chunk of text. For mean or sum pooling, we took the mean, or summed, across tokens within the current chunk of text.

### 4.4 Orthogonal Autocorrelated Sequences Model (*OASM*)

In general, brain responses nearby in time are more similar than brain responses further away in time. To model this temporal autocorrelation in brain responses, we construct a feature space for each dataset by (1) forming an *n*-dimensional identity matrix, where *n* is the total number of “stimuli” in the dataset for which distinct neural data were recorded (i.e., the number of rows in the original data matrix), and then (2) applying a Gaussian filter within blocks along the diagonal that correspond to temporally contiguous time points (i.e., within each passage in *Pereira2018*, each sentence in *Fedorenko2016*, and each story in *Blank2014*). This generates an autocorrelated sequence for each passage/sentence/story that is orthogonal to that of each other passage/sentence/story. Because of this orthogonality, OASM is by construction incapable of predicting brain responses when using contiguous splits.

We use the *scipy gaussian_filter1d* function to implement the Gaussian filter, and search through 48 sigma values (evenly spaced from 0.1 to 4.8) to find the optimal sigma value. We searched through 48 sigma values because *GPT2XL* has 48 layers, and so this allows for a fair comparison between the neural predictivity of both models.

It is important to clarify that although *OASM* is designed to model temporal autocorrelation, the neural variance it predicts is likely at least partly linguistically driven. This is because stimuli within a passage/sentence/story are also generally more linguistically similar than stimuli from different passages/sentences/stories. However, the key point is that in these datasets it is not possible to disentangle neural variance related to temporal autocorrelation from such linguistically driven neural variance and, furthermore, the representations of *OASM* yield essentially no theoretically interesting insight into human language processing.

### 4.5 Positional signals and word rate

Here we describe the construction of the positional signal and word rate (*PWR*) for each neural dataset.

#### Pereira2018

We originally constructed the positional model as a one-hot 4*D* vector, where each element corresponds to a given position of a sentence in the passage. We then applied a Gaussian filter to this one-hot vector in order to render activations for nearby positions more similar to one another. For this filter, we selected the *σ* value that maximized neural predictivity, searching through 48 (same number of layers as *GPT2XL*) *σ* values evenly spaced from 0 to 4.7. The word rate model was a 1*D* scalar which quantifies the number of words within each sentence.

#### Fedorenko2016

The primary finding in the paper which first reported this neural dataset, Fedorenko et al. [2016], was that neural activity exhibited ramping behavior across the words within each sentence. We thus first created a 1*D* positional signal whose activation increases across word positions within a sentence. We concatenated onto this 1*D* positional signal an 8*D* (because there were 8 words per sentence) one-hot positional vector to also model non-ramping neural activity. Because we expect positional representations in the brain to be more similar between adjacent words than between more distant words, we applied a Gaussian filter to the 8*D* positional signal. We searched across 48 *σ* values, evenly spaced from 0 to 4.7, and selected the *σ* value that maximized neural predictivity.

#### Blank2014

The positional model was a ramping signal which exhibited ramping behavior up to *N* TRs, and then displayed constant activation. For instance, when *N* = 4 the ramping signal was a vector of the following form: [0, 1, 2, 3, 3, …, 3]. For a given *N*, we also stacked on all ramping signals for smaller values of *N*, up to a minimum of *N* = 3. We selected the value of *N* which results in the highest test-set neural predictivity, and searched through values of *N* from 3 (corresponding to a 6s ramping signal) to 51 (corresponding to a 102s). The word rate model was a 1*D* scalar which quantified the number of words in the chunk of text corresponding to a given TR.

### 4.6 Syntax

Syntactic embeddings (*SYNTAX*) are derived by averaging together the embeddings of sentences that all share the same syntactic structure but differ in their content words. This procedure averages away differences in word/phrase/sentence-level meaning between the sentences, and keeps what is shared between them: syntactic structure (and abstract semantics associated with that structure). First, we create a word bank for each part-of-speech and dependency tag in the original sentences by passing 300, 000 sentences from the Generics KB corpus hug [2020] through the SpaCy transformer-based model [Honnibal and Montani, 2017]. Then, we run each sentence in *Pereira2018* through the SpaCy transformer-based model. We then generate new sentences, in which each content word from an original sentence is replaced with a content word of matching part-of-speech and dependency tag randomly sampled from the word bank. Each newly generated sentence is then once again passed through the SpaCy transformer-based model to get the token indices of the subtrees associated with each token; we ensure that, for each token index, the token indices of the subtree in the generated sentence match those in the original sentence. If not, we discard the generated sentence and try again. We continue until 170 new sentences with valid subtrees are generated for each original sentence. Finally, we take the cross-entropy loss of *GPT2XL* on each newly generated sentence and keep the 100 sentences with the smallest loss. To get a syntactic representation for each original sentence, we run each of these 100 new sentences through *GPT2XL* and average their representations. This method is highly similar to that of [Caucheteux et al., 2021], the main difference being that we keep function words from the original sentence. We keep function words for two reasons: (i) representations of function words have previously been used to generate representations of syntactic form for neural encoding Kauf et al. [2024], and (ii) coherent sentences were generated much more often when function words were left intact. Altogether, our syntactic representations combine the constraints on part-of-speech and dependency structure, as in Caucheteux et al. [2021], and the preservation only of function words, as in Kauf et al. [2024]. We used sum pooling to extract activations for *SYNTAX* because it resulted in higher neural predictivity than mean pooling or using the last token.

### 4.7 Regression methods

We used three linear regression methods throughout our paper. For all regression methods, we predict brain responses for each voxel/electrode/fROI independently. To compare our findings with Schrimpf et al. [2021], we employed ordinary least squares (OLS), which we implemented ourselves in order to run it on GPU. For all other analyses, we elected not to use OLS regression, because we are generally working with very ill-conditioned linear mappings due to high-dimensional feature spaces, relatively few samples compared to the number of features, and noisy targets. We instead used L2-regularized (ridge) regression for the majority of our analyses, which is the standard choice in the majority of neural encoding studies. When fitting regressions with both small and large feature spaces, we employed banded ridge regression to give small feature spaces their own L2 penalty [Dupré la Tour et al., 2022] **??**. This is because standard ridge regression assigns a single L2 penalty for all feature spaces, which can bias the regression against making use of small feature spaces (see Supplementary for ane expanded description of banded ridge regression). We fit all L2-regularized regression models with the *himalaya* package from Dupré la Tour et al. [2022]. We z-score all predictors across samples before training regressions, as is standard when using ridge regression in neural encoding studies.

#### Evaluation metrics

We define an encoding model (or simply model), *M*, as a set of feature spaces. To quantify the neural encoding performance of a model on the test set, we use both Pearson *r* and the out of sample *R*^2^ metric 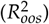 [Hawinkel et al.]. 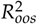 quantifies how much better a set of features performs at predicting held-out neural data compared to a model which simply predicts the mean of the training neural data (i.e. a regression with only an intercept term). Given predictions, 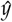, from a model *M* and predictions from a model with only an intercept term, denoted as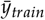, then:

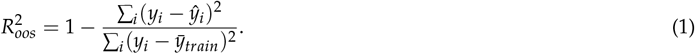

A positive (negative) value indicates that *M* was more (less) helpful than predicting the mean of training data. We elected to use 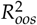 over the standard *R*^2^ because of this clear interpretation and because it is a less biased estimate of test set performance [Hawinkel et al.]. For variance partitioning analyses, we exclusively use 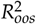 and not Pearson *r* because 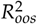 can be interpreted as the fraction of variance explained.

For both metrics, we take the mean neural predictivity across voxels/electrodes/fROIs for each participant, and compute the mean value across participants. We refer to 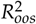 as *R*^2^ throughout the rest of the paper for brevity. For *R*^2^, we clip results to be at least 0 at the voxel/electrode/fROI level to prevent noisy values from biasing the mean downwards.

### 4.9 Train, validation, and test folds

For our main analyses, we constructed contiguous splits by ensuring brain responses from the same passage/sentence/story are not included in both train and test data. Due to low sample sizes, we employed a nested cross-validation procedure for each dataset. In the outer cross-validation loop, the brain responses were divided into training and testing sets. In the inner cross-validation loop, the training data was divided into training and validation sets to tune the L2 penalty values when using L2-regularized regression. To define the range of L2 penalty values, we generated a list of integers from *−*5 to 19, passed this list through numpy’s exp2 function, and then concatenated on 0. We describe the construction of our nested cross-validation procedure for each neural dataset below.

#### Pereira2018

During each outer fold, a single passage from each of the 24 semantic categories from one experiment was selected, and half of these passages were designated as the test set. By selecting at most a single passage from each semantic category, we ensured there were several passages in the training set which belonged to the same semantic category as the passages in the testing set. This equated to 8 test folds for experiment 1 (4 passages per semantic category, half selected for each loop) and 6 test folds for experiment 2 (3 passages per semantic category). During each inner fold, we again selected one passage from each semantic category, and half of these passages were designated as validation (leading to 7 inner folds for experiment 1, and 5 inner folds for experiment 2).

#### Fedorenko2016

For each outer fold, we selected 4 out of the 52 sentences as the test fold, resulting in 13 outer folds. For each inner fold, we once again selected 4 sentences as the validation set, resulting in 12 inner folds per outer fold.

#### Blank2014

For each outer fold, we selected a single story as the test fold, resulting in 8 outer folds. For each inner fold, each of the remaining stories served in turn as the validation set, resulting in 7 inner folds.

Because several previous studies using these datasets [Schrimpf et al., 2021, Kauf et al., 2024, Hosseini et al., 2024a, Oota et al., 2022, AlKhamissi et al., 2024, Aw et al., 2024, Hosseini et al., 2024b] employ shuffled splits, we also perform some analyses with shuffled splits. When using shuffled splits, data is arbitrarily placed into training and testing sets, such that data from the same passage/sentence/story can be in both the training and testing sets. To provide an example, this means that with the *Pereira2018* dataset it is possible for the first and third sentences within a passage to be in the training set, and the second sentence within the same passage to be in the testing set. In order to construct shuffled splits, we randomly shuffled the labels for each dataset that indicated which sentence belonged to which passage in *Pereira2018*, which word belonged to which sentence in *Fedorenko2016*, and which chunk of text corresponded to which story in *Blank2014*. We used numpy to perform this shuffling, and set the random seed to 42. Randomly shuffling the data labels made it so the testing and validation sets were no longer contiguous blocks of data, while still ensuring that the testing and validation sets were of the same size as the contiguous splits. We performed nested cross-validation in the exact same manner as described above when using shuffled splits.

When computing *R*^2^ across either inner folds or outer folds, we pooled predictions across folds and computed a single *R*^2^ as recommended by Hawinkel et al.. When computing Pearson *r*, we computed the correlation for each fold and then averaged across folds, as done in Schrimpf et al. [2021].

### 4.10 Selection of best layer

For each metric, we chose the layer that performs best across voxels/electrodes/fROIs on the test set with that metric. We used the test set to select the best layer to maintain consistency with Schrimpf et al. [2021]. For *Pereira2018*, we selected the best-performing layer for *EXP2* and *EXP3* separately. Due to the stochastic nature of untrained LLMs, we selected the best layer for 5 random seeds and report the average performance across seeds. When reporting the best layer, we refer to layer 0 as the input static layer, and layer 1 as the first intermediate layer. We selected hyperparameters for *OASM* and *PWR* for each metric separately, as well.

### 4.11 Percentage of LLM neural predictivity accounted for

Given a set of models *M*, we quantify the percentage of neural variance explained by an LLM that is accounted by *M* for a given participant through Eq. 3.

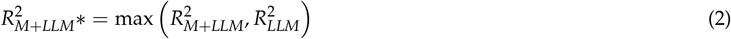

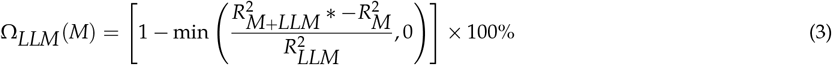

Here, *R*^2^ here denotes the mean *R*^2^ across voxels/electrodes/fROIs in a given participant. For a given participant, we first ensured that the neural predictivity of the *M+LLM* model was not lower than the predictivity of the *LLM* alone. Lower predictivity of *M+LLM* than *LLM* may happen due to overfitting, and in such cases we would unfairly state that *M* accounts for more of the neural predictivity of the *LLM* than it actually does. We denote this corrected neural predictivity as *R*_2*M*+*LLM*_*∗* (Equation 2). The numerator, *R*^2^_*M*+*LLM*_ *∗ − R*^2^_*M*_, quantifies the additional neural variance an *LLM* explains over *M* alone (this is the variance in neural predictivity of the LLM that is not explained by M). It is possible for this quantity to be less than 0 because adding the *LLM* to *M* may hurt neural predictivity (i.e., *R*^2^_*M*+*LLM*_ *∗ < R*^2^_*M*_). We then normalize this additional variance by the variance the *LLM* explains alone, *R*^2^_*LLM*_, and clip negative values since they are not interpretable. We subtract this quantity from 1 and multiply it by 100 to obtain the percentage of neural variance that the LLM explains that is also explained by *M*. We compute Ω for each participant separately, and a higher Ω value indicates that *M* accounts for more of the neural predictivity of the LLM.

### 4.12 Statistical testing

We perform two statistical tests within this study. When comparing the neural predictivity of an *LLM* to that of another model, we perform a one-side Wilcoxon signed-rank test across participants to determine whether the neural predictivity of the *LLM* is higher. The sample size for this test was *N* = 10 for *Pereira2018*, and *N* = 5 for *Fedorenko2016* and *Blank2014*. We exclusively perform this test on Pearson *r* values throughout the study.

We also perform a second statistical test to determine whether a regression that stacks together activations from a set of models, *M*, and from an *LLM* (*M+LLM)*, explains significantly more neural variance than *M* alone at the voxel/electrode/fROI level. When performing this test, we first selected the best sub-model within *M+LLM*, denoted as *M+LLM\**, and the best sub-model within *M*, denoted as *M\**, for each voxel/electrode/fROI separately. For instance, if comparing *PWR+GloVe+GPT2XL* to *PWR+GloVe*, we selected the sub-model which achieved the highest neural predictivity for each voxel/electrode/fROI for *PWR+GloVe+GPT2XL* and *PWR+GloVe*, separately. We then computed the squared error values on the test data for each of these models separately, which are obtained by subtracting the predictions of the regression model from the target brain responses and squaring this difference. We passed the squared error values from both models into a one-sided paired t-test which evaluates whether the squared error values from *M+LLM\** are significantly less than *M\**. The p-values were then FDR corrected Benjamini and Hochberg [1995] within each participant, and an *α* value of 0.05 is used to determine significance. The sample size for this t-test was *N* = 384 for voxels in *Pereira2018* with only data from *EXP2, N* = 243 for voxels in *Pereira2018* with only data in *EXP3*, and *N* = 627 for voxels with data in both experiments. For *Fedorenko2016*, the sample size was *N* = 416, and for *Blank2014* the sample size was *N* = 1317.

While squared error values are not always normally distributed, our sample sizes were large (the minimum sample size was 243), so we still opted to use a t-test over a non-parametric alternative Lumley et al. [2002]. We note that squared error values from a model are correlated, which means that the t-test is biased towards positive results which show that an LLM contributes significant variance over a set of simpler models. While this bias is not ideal, it is conservative in our case, making it harder to account for the neural variance that an LLM explains with simpler models.

## Footnotes

These across layer values are consistent with the code accompanying Schrimpf et al. [2021] and with Kauf et al. [2024]; they are not consistent the across layer pattern displayed in Figure 2c of Schrimpf et al. [2021] for reasons we are unaware of.

## References

Mariya Toneva and Leila Wehbe. Interpreting and improving natural-language processing (in machines) with natural language-processing (in the brain). Adv. Neural Inf. Process. Syst., pages 14928–14938, May 2019.

Martin Schrimpf, Idan Asher Blank, Greta Tuckute, Carina Kauf, Eghbal A Hosseini, Nancy Kanwisher, Joshua B Tenenbaum, and Evelina Fedorenko. The neural architecture of language: Integrative modeling converges on predictive processing. Proc. Natl. Acad. Sci. U. S. A., 118(45), November 2021.

Ariel Goldstein, Zaid Zada, Eliav Buchnik, Mariano Schain, Amy Price, Bobbi Aubrey, Samuel A Nastase, Amir Feder, Dotan Emanuel, Alon Cohen, and Others. Shared computational principles for language processing in humans and deep language models. Nat. Neurosci., 25(3):369–380, 2022.

Charlotte Caucheteux and Jean-Rémi King. Brains and algorithms partially converge in natural language processing. Commun Biol, 5(1):134, February 2022.

Richard Antonello, Aditya R Vaidya, and Alexander G Huth. Scaling laws for language encoding models in fMRI. Adv. Neural Inf. Process. Syst., abs/2305.11863, May 2023.

Alec Radford, Jeff Wu, R Child, D Luan, Dario Amodei, and I Sutskever. Language models are unsupervised multitask learners. 2019.

Eghbal A Hosseini, Martin Schrimpf, Yian Zhang, Samuel Bowman, Noga Zaslavsky, and Evelina Fedorenko. Artificial neural network language models predict human brain responses to language even after a developmentally realistic amount of training. Neurobiol Lang (Camb), 5(1):43–63, April 2024a.

Khai Loong Aw, Syrielle Montariol, Badr AlKhamissi, Martin Schrimpf, and Antoine Bosselut. Instruction-tuning aligns llms to the human brain. COLM, 2024.

Badr AlKhamissi, Greta Tuckute, Antoine Bosselut, and Martin Schrimpf. Brain-Like language processing via a shallow untrained multihead attention network. June 2024.

Jeffrey S Bowers, Gaurav Malhotra, Marin Dujmović, Milton Llera Montero, Christian Tsvetkov, Valerio Biscione, Guillermo Puebla, Federico Adolfi, John E Hummel, Rachel F Heaton, Benjamin D Evans, Jeffrey Mitchell, and Ryan Blything. Deep problems with neural network models of human vision. Behav. Brain Sci., 46:e385, December 2022.

Subba Reddy Oota, Jashn Arora, Veeral Agarwal, Mounika Marreddy, Manish Gupta, and Bapi Surampudi. Neural language taskonomy: Which NLP tasks are the most predictive of fMRI brain activity? In Marine Carpuat, Marie-Catherine de Marneffe, and Ivan Vladimir Meza Ruiz, editors, Proceedings of the 2022 Conference of the North American Chapter of the Association for Computational Linguistics: Human Language Technologies, pages 3220–3237, Seattle, United States, July 2022. Association for Computational Linguistics.

Carina Kauf, Greta Tuckute, Roger Levy, Jacob Andreas, and Evelina Fedorenko. Lexical-Semantic content, not syntactic structure, is the main contributor to ANN-Brain similarity of fMRI responses in the language network. Neurobiol Lang (Camb), 5(1):7–42, April 2024.

Eghbal Hosseini, Colton Casto, Noga Zaslavsky, Colin Conwell, Mark Richardson, and Evelina Fedorenko. Universality of representation in biological and artificial neural networks. bioRxiv, page 2024.12.26.629294, December 2024b.

Gavin Mischler, Yinghao Aaron Li, Stephan Bickel, Ashesh D. Mehta, and Nima Mesgarani. Contextual feature extraction hierarchies converge in large language models and the brain. Nature Machine Intelligence, 6(12):1467–1477, November 2024. ISSN 2522-5839. doi: 10.1038/s42256-024-00925-4. URL 10.1038/s42256-024-00925-4.

Francisco Pereira, Bin Lou, Brianna Pritchett, Samuel Ritter, Samuel J Gershman, Nancy Kanwisher, Matthew Botvinick, and Evelina Fedorenko. Toward a universal decoder of linguistic meaning from brain activation. Nat. Commun., 9(1):963, March 2018.

Evelina Fedorenko, Terri L Scott, Peter Brunner, William G Coon, Brianna Pritchett, Gerwin Schalk, and Nancy Kanwisher. Neural correlate of the construction of sentence meaning. Proc. Natl. Acad. Sci. U. S. A., 113(41):E6256–E6262, October 2016.

Idan Blank, Nancy Kanwisher, and Evelina Fedorenko. A functional dissociation between language and multiple-demand systems revealed in patterns of BOLD signal fluctuations. J. Neurophysiol., 112(5):1105–1118, September 2014.

Alexander G Huth, Wendy A de Heer, Thomas L Griffiths, Frédéric E Theunissen, and Jack L Gallant. Natural speech reveals the semantic maps that tile human cerebral cortex. Nature, 532(7600):453–458, April 2016.

Richard Antonello and Alexander Huth. Predictive coding or just feature discovery? an alternative account of why language models fit brain data. Neurobiol Lang (Camb), 5(1):64–79, April 2024.

Charlotte Caucheteux, Alexandre Gramfort, and Jean-Remi King. Disentangling syntax and semantics in the brain with deep networks. March 2021.

Charlotte Caucheteux, Alexandre Gramfort, and Jean-Rémi King. Evidence of a predictive coding hierarchy in the human brain listening to speech. Nat Hum Behav, 7(3):430–441, March 2023.

Zaid Zada, Ariel Goldstein, Sebastian Michelmann, Erez Simony, Amy Price, Liat Hasenfratz, Emily Barham, Asieh Zadbood, Werner Doyle, Daniel Friedman, Patricia Dugan, Lucia Melloni, Sasha Devore, Adeen Flinker, Orrin Devinsky, Samuel A Nastase, and Uri Hasson. A shared model-based linguistic space for transmitting our thoughts from brain to brain in natural conversations. Neuron, 112(18):3211–3222.e5, September 2024.

Ariel Goldstein, Eric Ham, Mariano Schain, Samuel Nastase, Zaid Zada, Avigail Dabush, Bobbi Aubrey, Harshvardhan Gazula, Amir Feder, Werner K Doyle, Sasha Devore, Patricia Dugan, Daniel Friedman, Roi Reichart, Michael Brenner, Avinatan Hassidim, Orrin Devinsky, Adeen Flinker, Omer Levy, and Uri Hasson. The temporal structure of language processing in the human brain corresponds to the layered hierarchy of deep language models, 2024. URL https://openreview.net/forum?id=95ObXevgHx.

Yoav Benjamini and Yosef Hochberg. Controlling the false discovery rate: A practical and powerful approach to multiple testing. J. R. Stat. Soc. Series B Stat. Methodol., 57(1):289–300, 1995.

Shailee Jain, Vy A Vo, Shivangi Mahto, Amanda LeBel, Javier Turek, and Alexander G Huth. Interpretable multi-timescale models for predicting fMRI responses to continuous natural speech. Adv. Neural Inf. Process. Syst., 33, October 2020.

Aniketh Janardhan Reddy and Leila Wehbe. Can fMRI reveal the representation of syntactic structure in the brain? Adv. Neural Inf. Process. Syst., 34:9843–9856, December 2021.

Amanda LeBel, Shailee Jain, and Alexander G Huth. Voxelwise encoding models show that cerebellar language representations are highly conceptual. J. Neurosci., 41(50):10341–10355, December 2021.

Wendy A de Heer, Alexander G Huth, Thomas L Griffiths, Jack L Gallant, and Frédéric E Theunissen. The hierarchical cortical organization of human speech processing. J. Neurosci., 37(27):6539–6557, July 2017.

Alexandre Pasquiou, Yair Lakretz, John Hale, Bertrand Thirion, and Christophe Pallier. Neural language models are not born equal to fit brain data, but training helps. July 2022.

Zhuoqiao Hong, Haocheng Wang, Zaid Zada, Harshvardhan Gazula, David Turner, Bobbi Aubrey, Leonard Niekerken, Werner Doyle, Sasha Devore, Patricia Dugan, Daniel Friedman, Orrin Devinsky, Adeen Flinker, Uri Hasson, Samuel A Nastase, and Ariel Goldstein. Scale matters: Large language models with billions (rather than millions) of parameters better match neural representations of natural language. October 2024. doi: 10.7554/elife.101204.1. URL 10.7554/eLife.101204.1.

Laurent Bonnasse-Gahot and Christophe Pallier. fmri predictors based on language models of increasing complexity recover brain left lateralization. In A. Globerson, L. Mackey, D. Belgrave, A. Fan, U. Paquet, J. Tomczak, and C. Zhang, editors, Advances in Neural Information Processing Systems, volume 37, pages 125231– 125263. Curran Associates, Inc., 2024. URL https://proceedings.neurips.cc/paper_files/paper/2024/file/e28e19d00b23fe0265f433fa05a96b06-Paper-Conference.pdf.

Amanda LeBel, Lauren Wagner, Shailee Jain, Aneesh Adhikari-Desai, Bhavin Gupta, Allyson Morgenthal, Jerry Tang, Lixiang Xu, and Alexander G Huth. A natural language fMRI dataset for voxelwise encoding models. Sci Data, 10(1):555, August 2023.

Rylan Schaeffer, Mikail Khona, and Ila Fiete. No free lunch from deep learning in neuroscience: A case study through models of the entorhinal-hippocampal circuit. NeurIPS, pages 16052–16067, August 2022.

Jeffrey S Bowers, Gaurav Malhotra, Federico Adolfi, Marin Dujmović, Milton L Montero, Valerio Biscione, Guillermo Puebla, John H Hummel, and Rachel F Heaton. On the importance of severely testing deep learning models of cognition. Cogn. Syst. Res., 82:101158, December 2023.

Olivia Guest and Andrea E Martin. On logical inference over brains, behaviour, and artificial neural networks. Computational Brain & Behavior, 6(2):213–227, June 2023.

Ansh Soni, Sudhanshu Srivastava, Konrad Kording, and Meenakshi Khosla. Conclusions about neural network to brain alignment are profoundly impacted by the similarity measure. bioRxiv, August 2024.

Colin Conwell, Jacob S Prince, Kendrick N Kay, George A Alvarez, and Talia Konkle. A large-scale examination of inductive biases shaping high-level visual representation in brains and machines. Nat. Commun., 15(1):9383, October 2024.

Evelina Fedorenko, Anna A Ivanova, and Tamar I Regev. The language network as a natural kind within the broader landscape of the human brain. Nat. Rev. Neurosci., 25(5):289–312, May 2024.

Richard Futrell, Edward Gibson, Harry J Tily, Idan Blank, Anastasia Vishnevetsky, Steven Piantadosi, and Evelina Fedorenko. The natural stories corpus. In Nicoletta Calzolari, Khalid Choukri, Christopher Cieri, Thierry Declerck, Sara Goggi, Koiti Hasida, Hitoshi Isahara, Bente Maegaard, Joseph Mariani, Hélène Mazo, Asuncion Moreno, Jan Odijk, Stelios Piperidis, and Takenobu Tokunaga, editors, Proceedings of the Eleventh International Conference on Language Resources and Evaluation (LREC 2018), Miyazaki, Japan, May 2018. European Language Resources Association (ELRA). Genericskb: A knowledge base of generic statements. Allen Institute for AI, 2020.

Matthew Honnibal and Ines Montani. spaCy 2: Natural language understanding with Bloom embeddings, convolutional neural networks and incremental parsing. To appear, 2017.

Tom Dupré la Tour, Michael Eickenberg, Anwar O Nunez-Elizalde, and Jack L Gallant. Feature-space selection with banded ridge regression. Neuroimage, 264:119728, December 2022.

Stijn Hawinkel, Willem Waegeman, and Steven Maere. Out-of-Sample r2: Estimation and inference. Am. Stat., pages 1–11.

Thomas Lumley, Paula Diehr, Scott Emerson, and Lu Chen. The importance of the normality assumption in large public health data sets. Annu. Rev. Public Health, 23:151–169, 2002.

