## Extended Data Figures for "Illusions of Alignment Between Large Language Models and Brains Emerge From Fragile Methods and Overlooked Confounds"

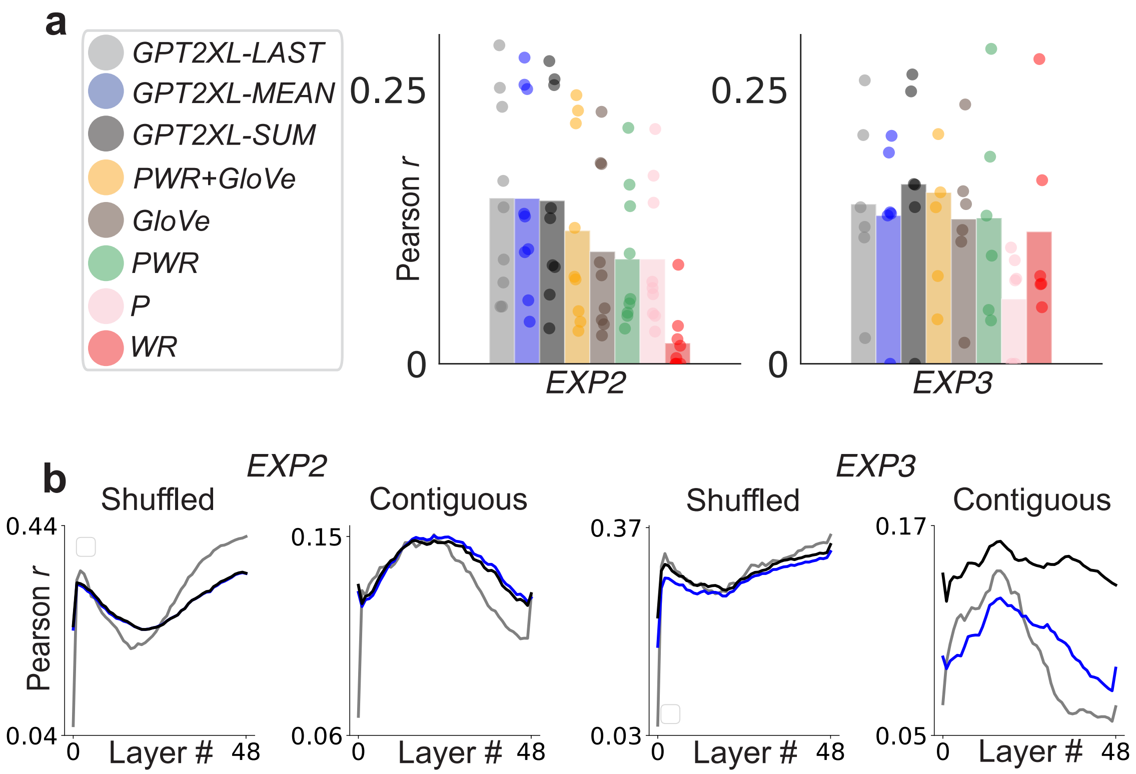


Figure 1: **a)** Neural predictivity of models for each sub-experiment within *Pereira2018*. The *PWR* model is further divided into its sub-models: *position* (*P*) and *word rate* (WR). **b)** *GPT2XL* across-layer neural predictivity for each experiment in *Pereira2018*

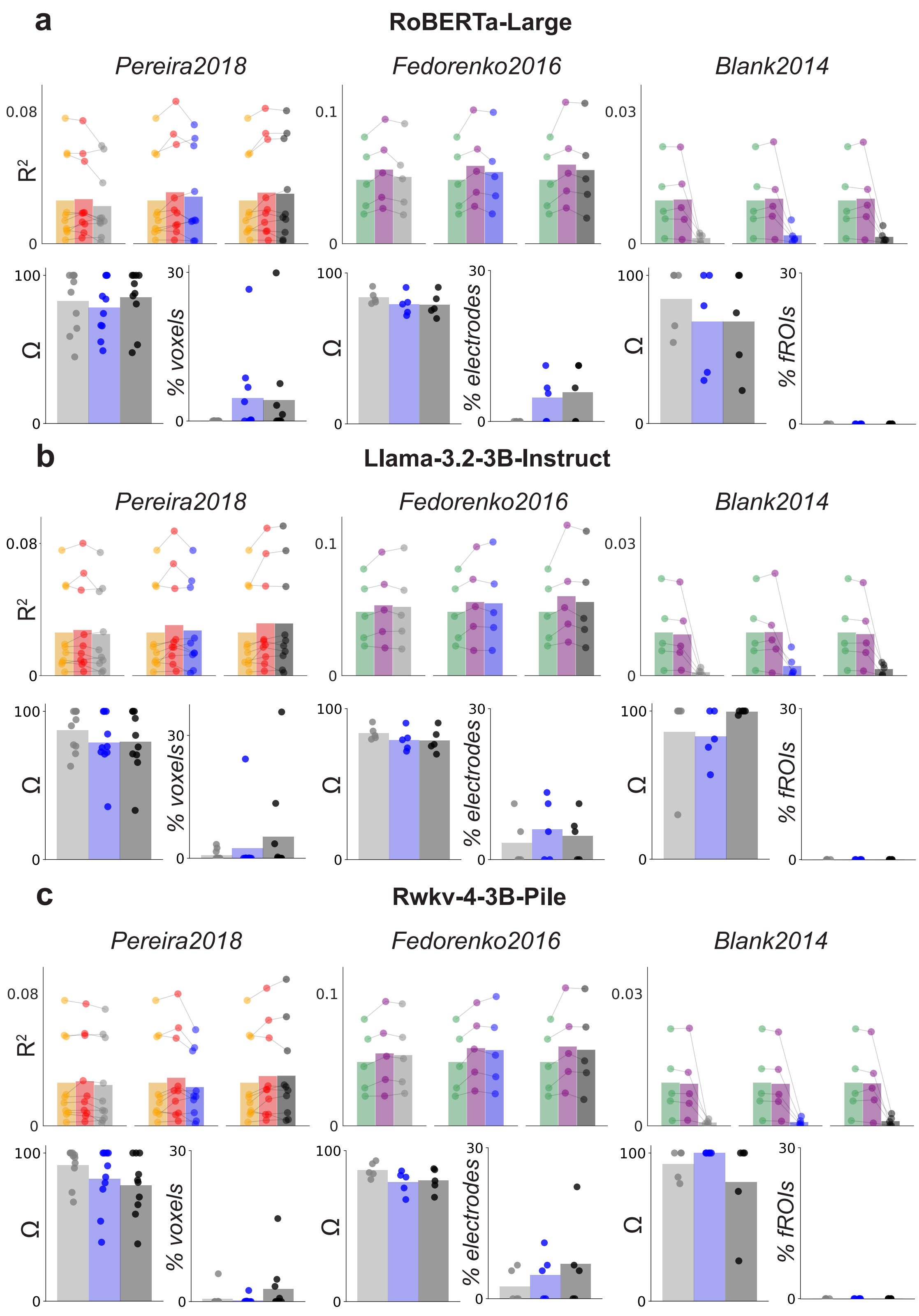


Figure 2: Color scheme is the same as in Figure 5, except that *GPT2XL* colors now correspond to appropriate LLM. All sub-panels are identical in style to Figure 5*c* and *d*. **a)** Results for *RoBERTa-Large*, **b)** Results for *Llama-3.2-3B-Instruct*, **c)** Results for *Rwkv-4-3B-Pile*.

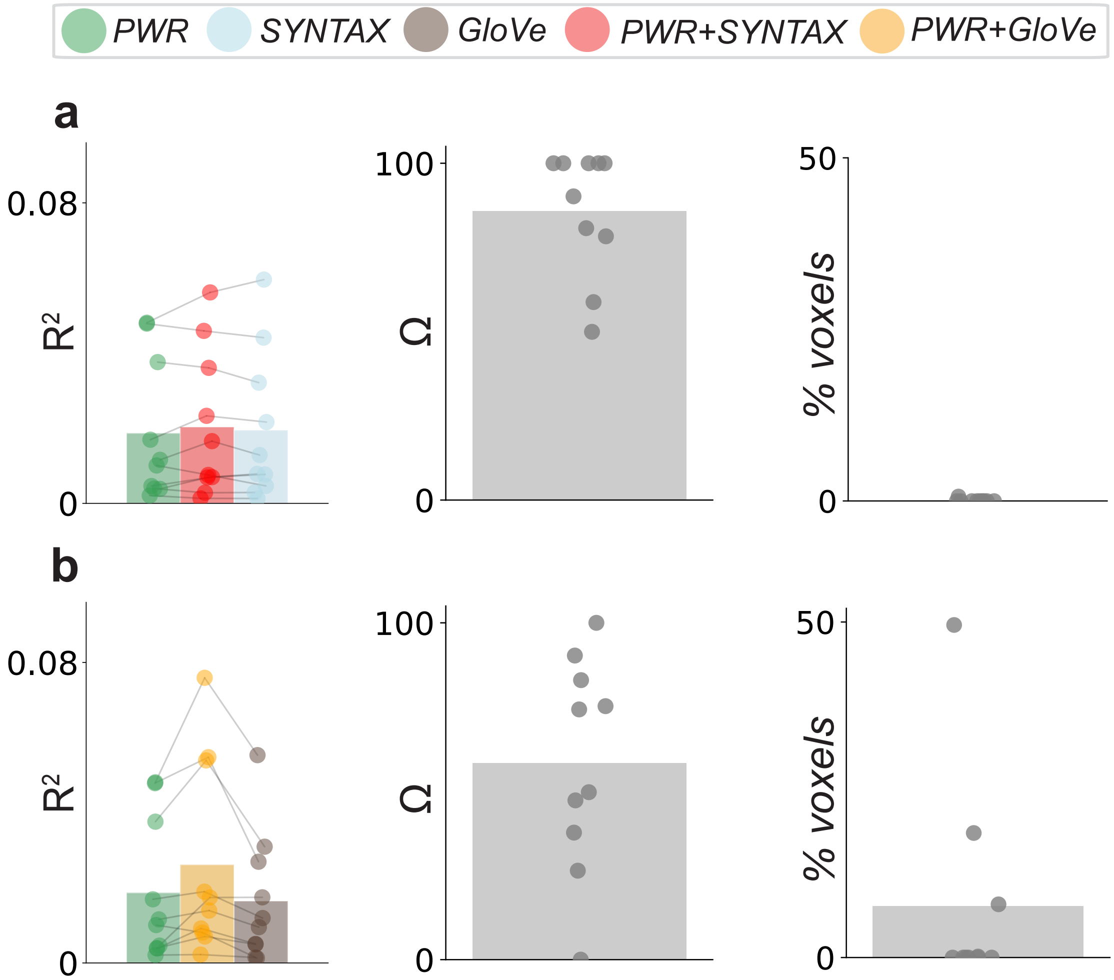


Figure 3: **a)** Right-side shows neural predictivity for *PWR*, *PWR+SYNTAX*, and *SYNTAX*. Middle shows the percentage of *SYNTAX* neural predictivity that *PWR* accounts for, or $\Omega_{\text{SYNTAX}}(PWR)$. Left side shows the percentage of voxels where *PWR+SYNTAX* explains more neural variance than *PWR* alone. **b)** Same as **(a)**, except the *SYNTAX* model is replaced with *GloVe*. Each dot in bar plot shows values for a given participant.

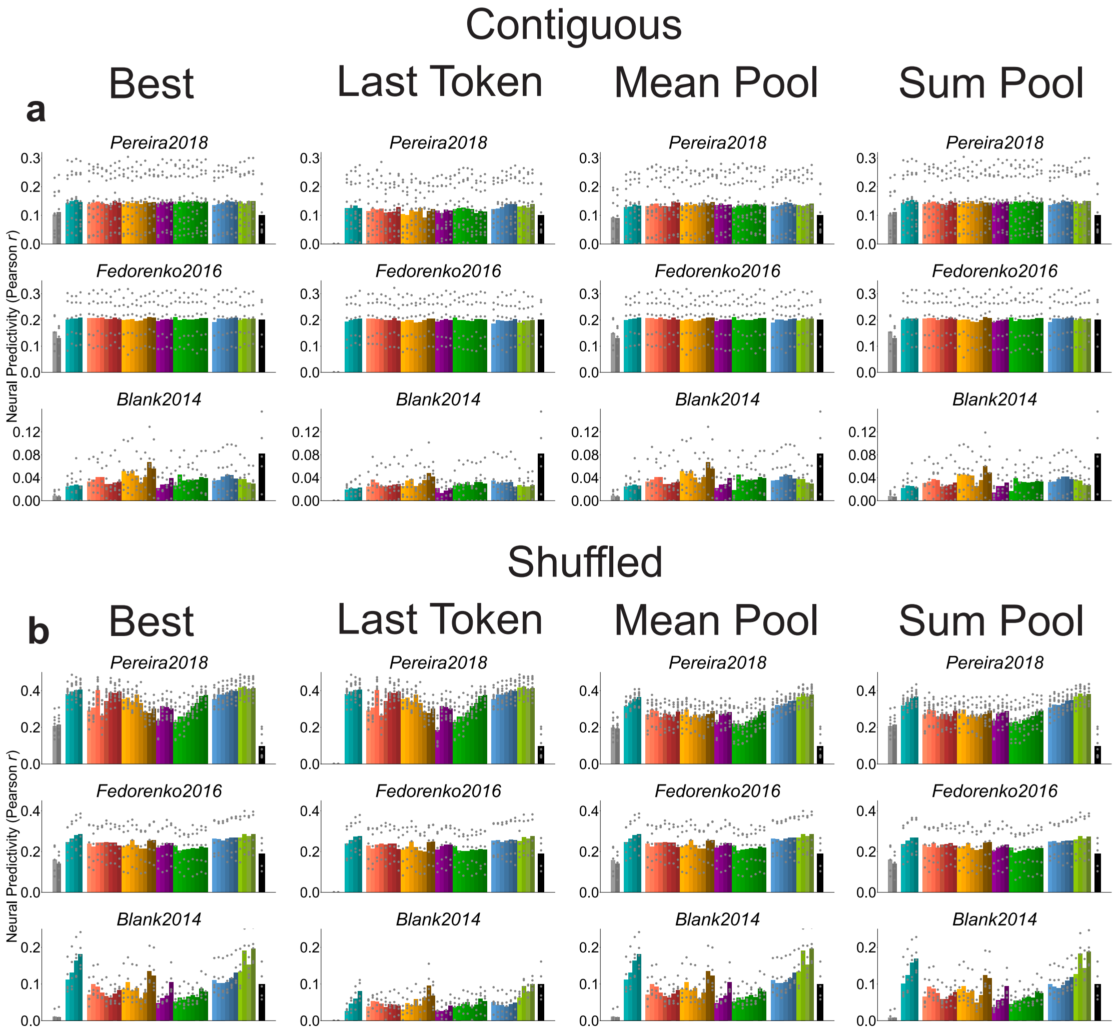


Figure 4: **a)** Model comparisons of neural predictivity when using contiguous splits, showing the neural predictivity for each individual participant (gray dots). **b)** Same as **(a)** but for shuffled splits.

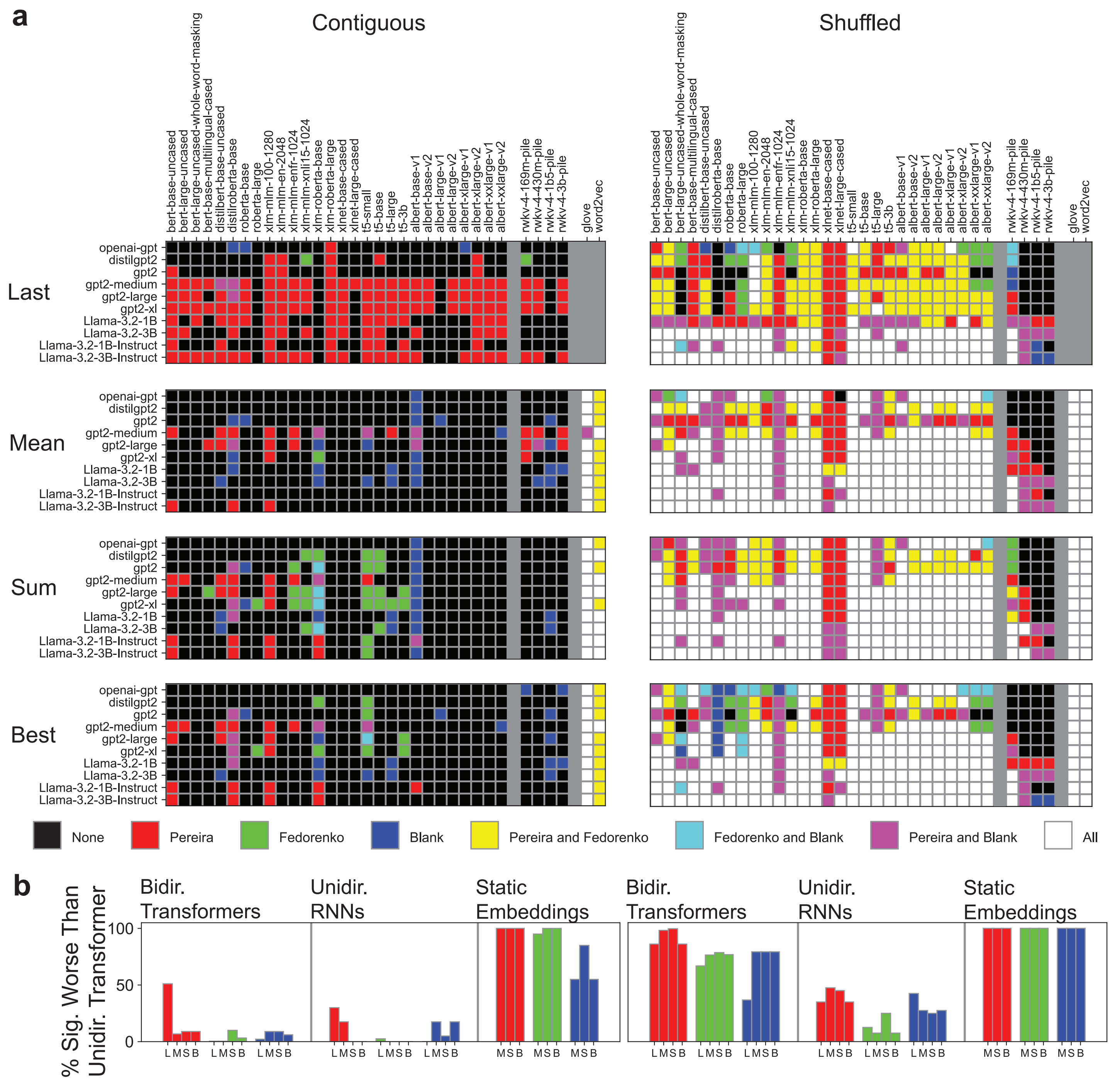


Figure 5: **a)** Significance results (one-way Wilcoxon signed-rank test, $p < 0.05$) depicting when each unidirectional transformer model (vertical axis) significantly outperforms each model of each other model class (horizontal axis). Gray bars separate model classes (bidirectional transformers, unidirectional RNNs, and static embeddings). The color of each box depicts what combination of datasets the comparison is significant for. Note that results are omitted for comparisons with static embeddings in the last-token case, as this activation extraction method is not appropriate for non-contextual models. **b)** The percentage of significance tests for which a unidirectional transformer significantly outperforms a model for each other class of models. As in **(a)**, red, green and blue correspond to *Pereira2018*, *Fedorenko2016*, and *Blank2014*, respectively. Adjacent bars correspond to the different feature extraction methods, with L, M, S and B denoting last-token, mean-pool, sum-pool, and best, respectively.
